# Design of amyloidogenic peptide traps

**DOI:** 10.1101/2023.01.13.523785

**Authors:** Danny D. Sahtoe, Ewa A. Andrzejewska, Hannah L. Han, Enrico Rennella, Matthias M. Schneider, Georg Meisl, Maggie Ahlrichs, Justin Decarreau, Hannah Nguyen, Alex Kang, Paul Levine, Mila Lamb, Xinting Li, Asim K. Bera, Lewis E. Kay, Tuomas P.J. Knowles, David Baker

## Abstract

Segments of proteins with β-strand propensity can self associate to form amyloid fibrils associated with many diseases. These regions often adopt alternative structures in their folded states, or are intrinsically disordered in solution, making it difficult to generate binders or inhibitors with existing strategies. Here we describe a general approach to bind such segments in β-strand and β-hairpin conformations using *de novo* designed scaffolds that contain deep peptide binding clefts flanked by β-strands that form hydrogen bonds to the peptide upon binding. The designs bind their cognate peptides *in vitro* with nanomolar affinities and in mammalian cells. The crystal structure of a designed protein-peptide complex is close to the design model, and NMR characterization reveals how the peptide binding cleft is protected in the apo state. We use the approach to design binders to segments of the amyloid forming proteins Transthyretin, Tau, Serum amyloid A1 and Aβ42. The Aβ binders block assembly of Aβ fibrils as effectively as the most potent of the clinically tested antibodies to date.

## Introduction

Many proteins contain segments that only become ordered upon binding a target (Tsai, Xu, and Nussinov 1998; Wright and Dyson 2009; Shammas et al. 2016). A particularly interesting example of such disorder-to-order transitions are amyloidogenic sequences found in proteins such as Aβ42, Tau and Serum amyloid A1. These regions can aggregate into amyloid fibrils via strand-strand interactions and are associated with amyloidosis and associated diseases both inside and outside the central nervous system (G.-F. Chen et al. 2017; Gamblin et al. 2003; Lu et al. 2014; Iakovleva et al. 2021; Knowles, Vendruscolo, and Dobson 2014; Chiti and Dobson 2006). Although the correlation between amyloid formation and neurodegenerative disease remains incompletely understood, designed binders to amyloid forming segments of these proteins could have utility both as diagnostics and therapeutics. However, it is difficult to raise antibodies against the monomeric form of amyloid due to their strong tendency for self association; this also complicates the systematic generation of binders using library selection methods, although some molecules have been evolved through these methods (Linse et al. 2020; Boutajangout et al. 2019; Panza et al. 2019). While there have been considerable advances in computational protein design, the multiplicity of conformations complicates design of binders to disordered protein segments, and the computational design of binders to amyloid forming segments of proteins remains an outstanding challenge.

We reasoned that this challenge could be overcome by taking advantage of the β-strand forming propensity of amyloidogenic peptides. Binding of peptides in β-strand conformation has been observed in nature (Remaut and Waksman 2006; Watkins and Arora 2014), and the regularity of the β secondary structure has been exploited to computationally design interactions between pairs of folded proteins such as homodimers, binders against natural target proteins with exposed β-strands, as well as nanoscale multi-subunit hetero-oligomers (Stranges et al. 2011; Lin et al. 2017; Sahtoe et al. 2021, 2022). To design binders to peptides in extended β-strand conformations, we sought to create scaffolds that could provide β-strand pairing interactions to all the backbone amide and carbonyl atoms of the peptide such that the peptide strand complements a β-sheet on the scaffold (Fig 1A).

**Figure 1.**
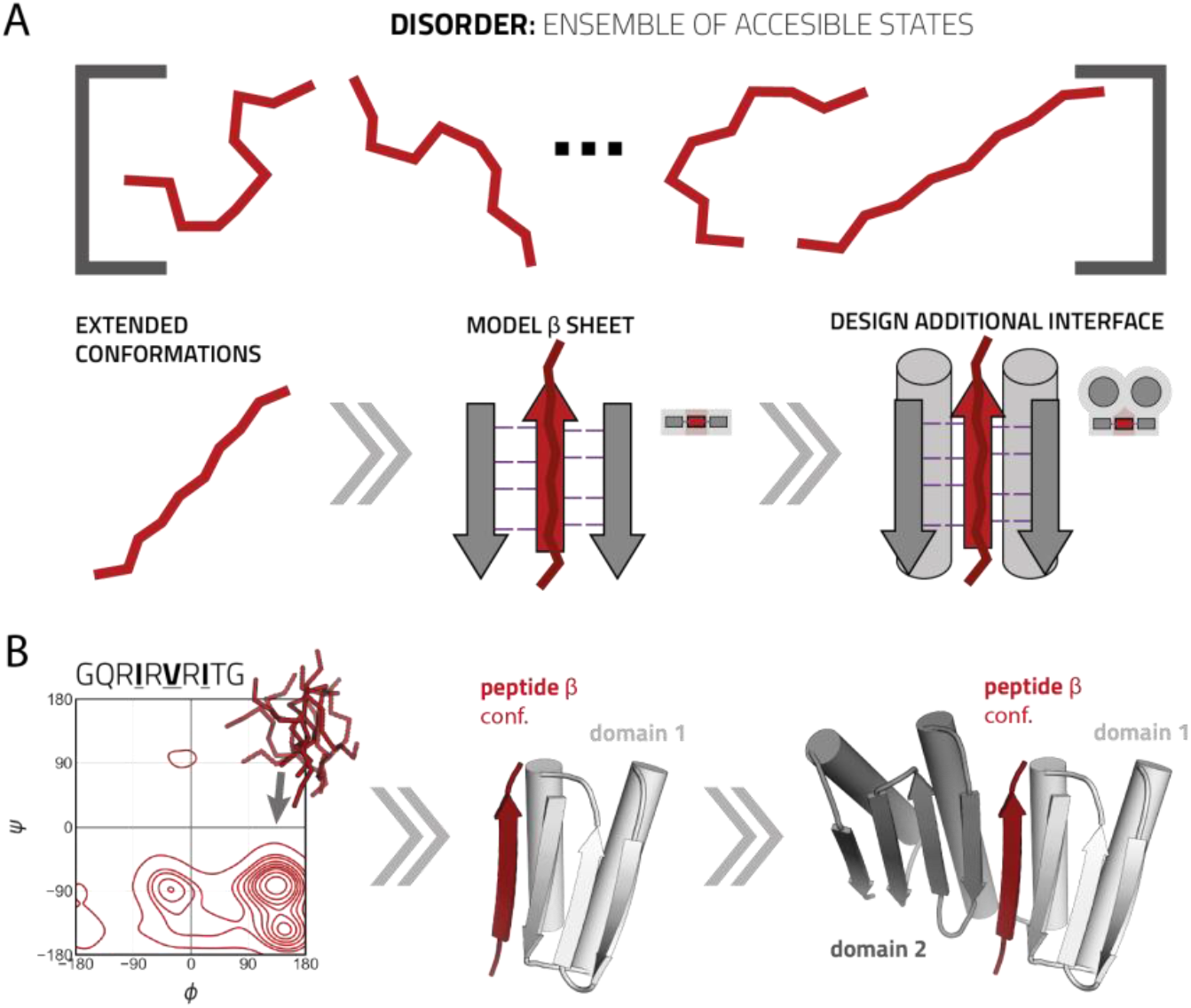
Design approach for binding disordered protein fragments. **a:** Intrinsically disordered regions of proteins and peptides have large conformational freedom but may be forced into predefined conformations such as β-strands that can be efficiently targeted using strand-strand interactions. **b:** Molecular mechanics simulation of a model peptide (red) shows it adopts a wide range of conformations (left) but can be modeled in a β-conformation while strand-pairing to a de novo protein (light gray, middle). A second domain (dark gray, right) can be designed that provides strand-strand interactions to the other side of the peptide creating a single chain protein with a deep and complementary peptide binding cleft.

Starting from Fold-It designed proteins with mixed α/β topology (Koepnick et al. 2019), we designed additional strands and helices to create scaffolds with a single central β-strand missing from an extended β-sheet. The sheet is buttressed by α-helices which pack on one another to support the structure in the absence of the bound peptide (see Fig 1b and s1 and Methods). Rosetta combinatorial sequence design calculations were then used to optimize the sequences of both the scaffold and the peptide for high affinity binding (we reasoned that such “two-sided” designs would be an easier starting point than “one sided” designs against amyloid forming peptides where only the sequence of the binders are allowed to be optimized). Designs with favorable interaction energy, few unsatisfied buried polar atoms and high shape complementarity, and for which Rosetta structure predictions were close to the designed scaffold and complex structures were selected for experimental characterization.

The selected designs without their cognate peptides were encoded in synthetic genes with an N-terminal polyhistidine affinity tag, expressed in *Escherichia coli*, and purified using immobilized nickel affinity chromatography (IMAC) followed by size exclusion chromatography (SEC). Despite the absence of the peptide, a large number of designs expressed well and were monodisperse in SEC. Bicistronic vectors were generated for each of the monodisperse designs; the first cistron encodes sfGFP fused at its C-terminus to the designed peptide, and the second cistron the polyhistidine tagged designed binder. After expression of the bicistronic constructs, binding of the GFP-peptide fusion to the his-tagged binder was assessed by SDS-PAGE following purification by IMAC and SEC.

The binding of six designs (figure 2a and figure s2 and table s1 and supplementary spreadsheet) that were well expressed, soluble, and monodisperse by SEC, to their designed peptide targets was further characterized using biolayer interferometry (BLI) by immobilizing chemically synthesized biotinylated peptides on streptavidin sensors and dipping these into a solution with the designed binding partner. The interaction kinetics ranged from 10^4^ M^-1^ s^-1^ to 10^2^ M^-1^ s^-1^ for association and between 0.17 s^-1^ - 10^−4^ s^-1^ for the dissociation (table s2). The equilibrium dissociation constant *K*_D_ ranged from 44 μM to 150 nM with no clear distinction between single strand and hairpin binders (figure 2a and table s2). For design C104 we confirmed binding in an orthogonal SEC binding assay (figure s3a).

**Figure 2.**
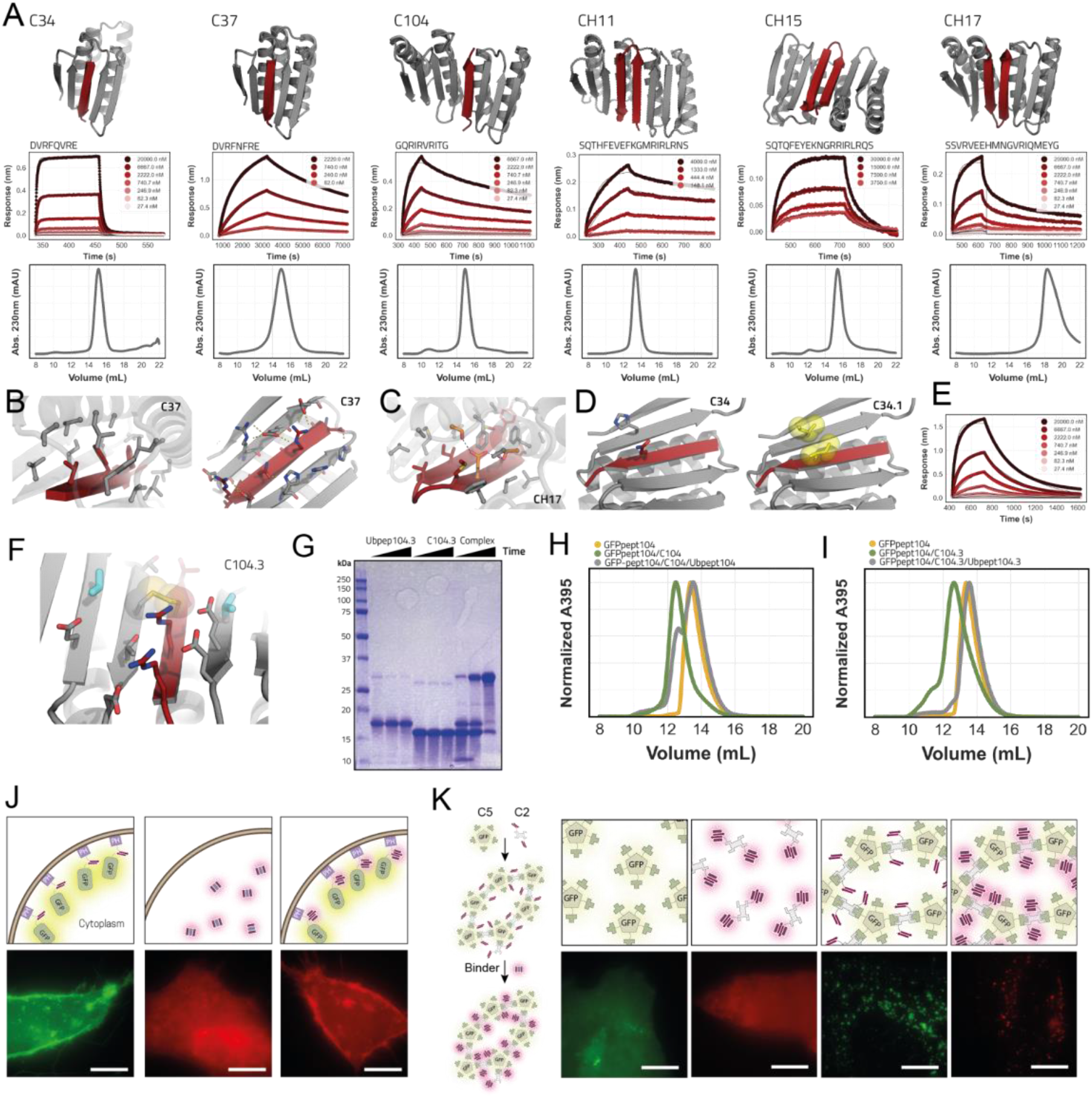
Characterization of designed peptide binders. **a:** Designed models for peptide binders (binder in gray, peptide in dark red). Respective BLI traces with kinetic fits and SEC (S75 increase 10/300) chromatograms of the binders are shown below the models. **b:** Detailed views of the solvent exposed interface of C37 (right) and the buried interface (left). C-alpha atoms as spheres. **c:** Detailed view of the buried part of the interface of hairpin binder CH17 with the designed hydrogen bond network depicted in orange sticks. **d:** Models of parent design C34 (left) and C34.1 (right) where an hydrophobic interaction pair (yellow sticks/spheres) is introduced to improve affinity. **e:** BLI trace of C34.1 binding its peptide that is immobilized on the biosensors. **f:** View of the designed interface disulfide on C104.3 (disulfide in spheres and sticks; additional redesigned residues in cyan). **g:** Non reducing SDS-PAGE gel showing disulfide formation (timepoints; t=0, t=90min t=overnight). **h:** SEC trace of preformed non-covalent C104 complex + GFP-pep104. **i:** SEC trace of preformed covalent disulfide linked C104.3 complex + GFP-pep104. **j:** Fluorescent microscopy images of mScartlet CH15.1 localization to membranes in HeLa cells. Scale bars 10 μm **k:** Fluorescent microscopy images of mScartlet CH15.1 localizing to designed intracellular GFP positive protein punctae in HeLa cells. Scale bars 10 μm.

Single amino acid substitution of the buried residue Val6 in the peptide of C104 to Arg completely disrupted binding in BLI suggesting that the designed binding mode is recapitulated (figure s3b and s3c).

The designed peptides are amphipathic with an alternating hydrophilic-hydrophobic side chain pattern (figure s3d). Beyond the backbone β-strand hydrogen bonding, the peptide-binder interaction consists of somewhat separable solvent exposed and solvent shielded interfaces. The solvent inaccessible part of the interface consists primarily of the hydrophobic residues that closely pack against the hydrophobic core of the binder and drive the association between peptide and binder (figure 2b, left). In design CH17 these interactions are accompanied by designed buried hydrogen bond networks (figure 2c) (Boyken et al. 2016). The solvent exposed portion of the interface (figure 2b, right) is composed primarily of salt bridges and hydrogen bonds that likely make less of a contribution to the overall interface energy because of competition with water. Because the hydrophobic-hydrophilic patterning is shared among the designed peptides, not all designs are able to fully discriminate between cognate and non-cognate peptides enabling them to sequester a broad range of peptides that have similar physicochemical properties (figure s4). Design CH17 that contains buried hydrogen bond network is however more selective to its cognate peptide, because binding of a non-complementary peptide would bury polar residues that are not satisfied with a hydrogen bond donor/acceptor (figure s4) disfavoring binding.

We explored the possibility of increasing peptide binding affinity by introducing hydrophobic interaction pairs across solvent exposed parts of the interface using combinatorial side chain design in Rosetta. Introduction of an exposed hydrophobic interaction pair in design C34.1 improved the *K*_D_ 6-fold to 2 μM from 12 μM in parent design C34 (figure 2d-e and table s2). In CH15.1 we introduced 3 hydrophobic interaction pairs that when combined led to a 400-fold improvement of the *K*_D_ from 40 μM to 100 nM compared to the parent CH15 design (figure 3f and s5a-b and table s2). The modified designs remained monomeric, indicating that these surface substitutions are generally well tolerated (figure s5c-d).

**Figure 3.**
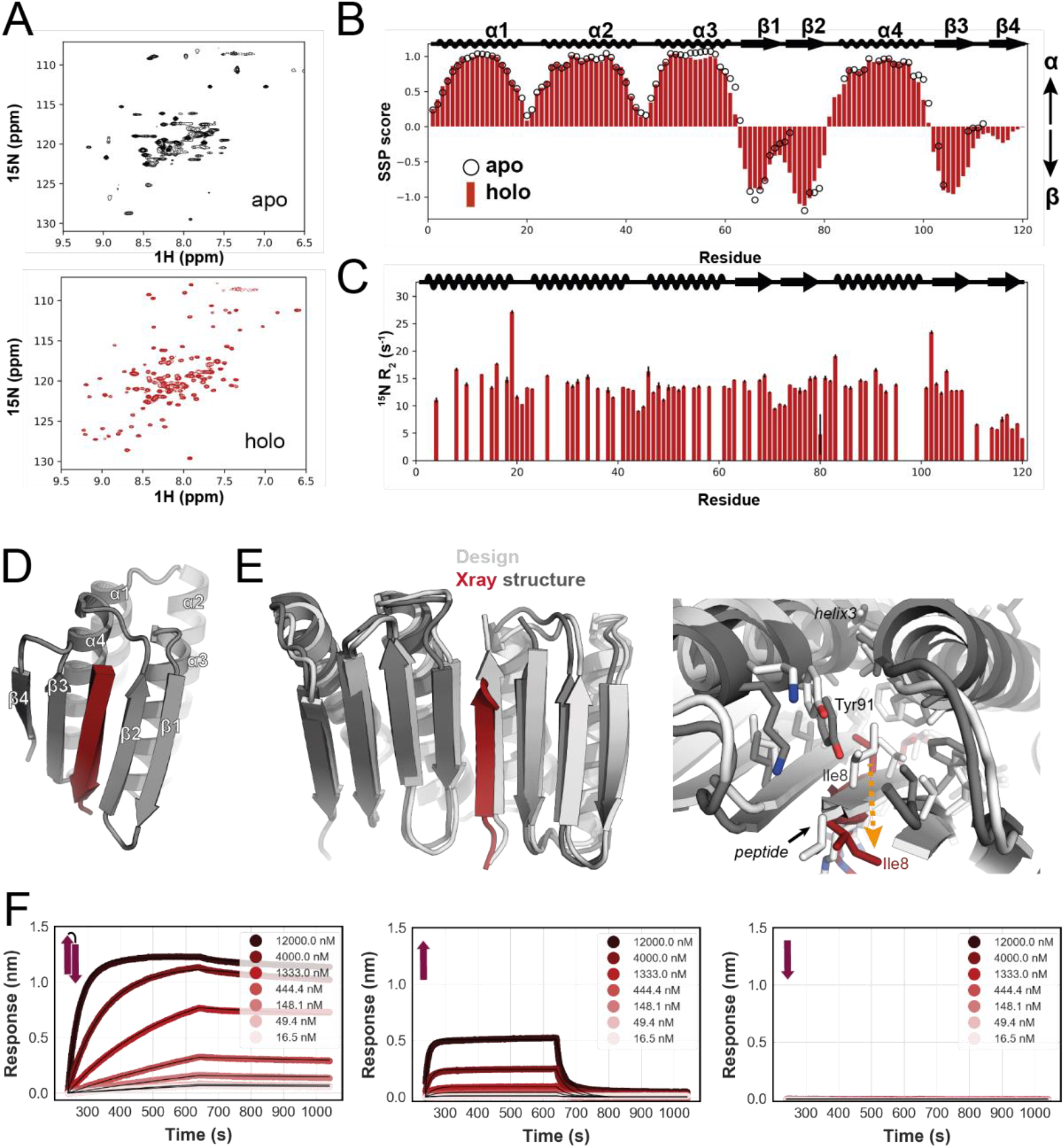
Structural characterization of designs. **a:** NMR spectra of ^15^N labeled C34 in absence (top) and in presence of 10-fold excess target peptide (bottom), 25°C. **b:** Secondary structure propensity as a function of residue, based on backbone ^1^H, ^13^C, and ^15^N chemical shifts recorded at 50°C using the SSP program (Marsh et al. 2006). SSP scores for the apo-form are shown with open circles, while those for the peptide-bound state are indicated with bars. The putative secondary structure of the designed protein is indicated above the plot. Positive values of SSP indicate α-helical structure, while negative values denote β-strands. **c:** ^15^N transverse relaxation rates as a function of residue. Low values, such as those in putative β4, indicate rapid time-scale dynamics, and are consistent with poorly formed structure. **d:** Designed model of C34. **e:** On the left; Overlay of the design model of a surface redesigned version of C104 (gray) and the crystal structure (colors). On the right; Detailed interface view of design (gray) and crystal structure (colors) with Ile8 shift indicated with orange dotted arrow. **f:** Binding of CH15.1 to its hairpin peptide (left), the individual N-terminal strand (middle) and C-terminal strand (right) of the hairpin in BLI.

Disulfide functionalization could enable redox control of binding activity for a variety of biotechnological applications. We searched for positions that could host a disulfide bridge across the interface of C104 using the Disulfidize mover in Rosetta (Fleishman et al. 2011; Bhardwaj et al. 2016) and found several positions where low energy disulfides could be modeled (figure 2f and figure s6a). For designs C104.2 and C104.3 we confirmed through non-reducing SDS-PAGE analysis that disulfides indeed formed (figure 2g and figure s6a-b). For C104.3 this result was further validated in a SEC subunit exchange experiment where we first reconstituted the non-covalent complex between C104 and its peptide fused to the c-terminus of ubiquitin, as well as the disulfide linked complex between C104.3 and its cysteine containing peptide fused to c-terminus ubiquitin. When the preformed non-covalent complex was mixed with GFP-104 and ran over SEC, GFP-104 co-eluted with C104 as observed through the absorbance at 395 nm indicating GFP-104 could exchange with ubiquitin-peptide fusion to bind C104 (figure 2h). In contrast, the peptide in the covalent C104.3 complex could not be outcompeted when it was mixed with GFP-P104 due to the disulfide bridge (figure 2i and figure s6c).

To examine the functionality of the designs in mammalian cells, we transfected HeLa cells with a construct where the peptide of CH15.1 was fused to the N-terminus of GFP and to the C-terminus of phospholipase-C Pleckstrin Homology domain that binds Phosphatidylinositol 4,5-bisphosphate at the outer plasma membrane (Várnai and Balla 1998). Fluorescent microscopy analysis showed that the plasma membrane of transfected cells were labeled green. When cells were also transfected with mScarlet labeled CH15.1 binder, GFP and mScarlet colocalized at the plasma membrane indicating binding (figure 2j and figure s7a-b). In control cells that were transfected with just mScarlet-CH15.1, or with mScarlet-CH15.1 and a mutant peptide intended to disrupt binding, no colocalization was observed indicating the interaction takes place through the designed interface (figure 2j and figure s7c).

In a second cell based experiment we tested whether the binder-peptide interaction could localize to intracellular 2-component protein puncta. The first component is a homopentamer fused to GFP and one half of a designed LHD heterodimer (Sahtoe et al. 2022) whereas the second component is a pseudo-C2 symmetric design that presents two copies of the other half of the designed heterodimer, and is also fused to the peptide of CH15.1 (figure 2k). When the homopentamer was expressed in HeLa cells we observed a diffuse GFP distribution. Upon co-expression of the second component a protein network was formed through the designed LHD heterodimer interfaces as observed by the formation of GFP puncta (figure 2k). Whenever mScarlet tagged CH15.1 binder was also present it was recruited to the puncta whereas in control experiments where the puncta did not form or where the mutant peptide was transfected, mScarlet CH15.1 binder was distributed uniformly throughout the cell indicating the peptide-binder pair can specifically associate within the crowded environment of the cell (figure 2k and figure s7d-f).

Small peptides are useful as affinity tags to bind and localize tagged protein partners into larger molecular assemblies. In nature, this method of protein recruitment is commonly used to regulate various cellular processes in a dynamic fashion. To demonstrate the utility of our designs towards such applications and also for use in novel customizable protein materials, we rigidly fused binder C37 to the LHD284B9 component of the LHD hetero-oligomer system that consist of de novo designed protein building blocks that can be assembled into a large variety of multiprotein complexes (Sahtoe et al. 2022). Fusion creates single chain proteins with two different interfaces; one peptide binding interface and one LHD heterodimer interface. Mixing for instance GFP tagged peptide of C37 with C37LHD284B9 creates a heterodimer. This assembly can further be expanded by the addition of for example LHD284A82 creating a heterotrimer. We confirmed the assembly of this complex via SEC (figure s8).

In the absence of peptide the binder contains a vacant cleft which exposes a hydrophobic core. Structure prediction methods predict that this cleft closes to form a continuous sheet in the apo state suggesting the designs are structurally dynamic (figure s9a). To study this we recorded an ^15^N,^1^H-HSQC NMR spectrum of unbound C34. The spectrum showed broadened resonances (figure 3A top), suggesting the occurrence of exchange processes on the millisecond-timescale. While the observed conformational dynamics is noteworthy and is the subject of further investigation, it prevents a straightforward structural characterization for most of the designed β-strand regions in the absence of the peptide. As a result large portions of putative strand β3 and the whole of putative strand β4 could not be assigned and therefore the secondary structure propensities (open circles in figure 3B and figure s9b) (Marsh et al. 2006) could not be calculated for residues within these regions, although it is clear that well-defined, stable structure is absent. This contrasts with the structure predictions for apo C34 in which the entire sheet is expected to form in the absence of peptide (figure s9a). For the rest of the protein, however, resonance assignments could be obtained at 50°C, where the high temperature decreases the effect of exchange, and the secondary structure propensity is close to the design even in the absence of the peptide (figures 3B and S9b,c).

In contrast to the free state, the NMR spectrum of the bound state shows sharp signals (figure 3A bottom), indicating that the exchange process is quenched in the presence of the peptide. The secondary structure is as designed (figure 3b, blue bars, and figure s9c), except for β4 which has lower β strand propensity, as confirmed by ^15^N transverse relaxation (R_2_) experiments that indicate an increase in fast time-scale dynamics in this region (figure 3C). We confirmed that the peptide binds in the designed orientation by measuring intermolecular NOE contacts between it and C34 (figure s9d).

We obtained a 2.3 Å resolution crystal structure of a variant of C104, C104.1, where all the surface residues outside the interface were redesigned using ProteinMPNN (Dauparas et al. 2022). The crystal structure recapitulates the designed model with both individual domains clamping the peptide in a β-strand conformation (figure 3e and table s2). The individual domains superimpose well with the design model. The majority of the peptide is resolved in the electron density and binds in a β-strand conformation with the apolar residues buried in the designed cleft (figure 3e and figure s9e). A deviation from the designed model at helix3 shifts Tyr91 towards the peptide binding pocket in the crystal structure partially occluding it (figure 3e). As a result peptide residue Ile8 is displaced (figure s9f) and the last few residues of the peptide are disordered in the crystal and not modeled (see methods).

While we were not able to obtain a crystal structure of a hairpin binding design, strand deletion experiments support that these peptides bind to the scaffold in a hairpin conformation rather than through single strand insertion: the binding of each individual strand of the CH15.1 hairpin to the CH15.1 binder is weaker than binding of the whole hairpin by BLI (figure 3f).

Encouraged by the biochemical and structural validation of our design approach on the two sided binder design challenge, we next investigated whether the approach could generate binders to naturally occurring peptide or protein segments which form amyloids in a range of disease states. This is a more challenging “one sided” design problem because the target sequence is fixed. Amyloid fibril deposits can form in the central nervous system as is the case for Aβ42, Microtubule associated protein Tau, and alpha-synuclein but also extra-cerebrally like in transthyretin and serum amyloid A1 mediated amyloidosis (Lu et al. 2014; Bloom 2014; Muchtar et al. 2021). The fibrils form through strand-strand mediated oligomerization/fibrillization and are harmful to cells and tissues (Knowles, Vendruscolo, and Dobson 2014; Chiti and Dobson 2006). We aimed to design binders to fibril forming regions to block or modulate fibril assembly (figure 4a).

**Figure 4.**
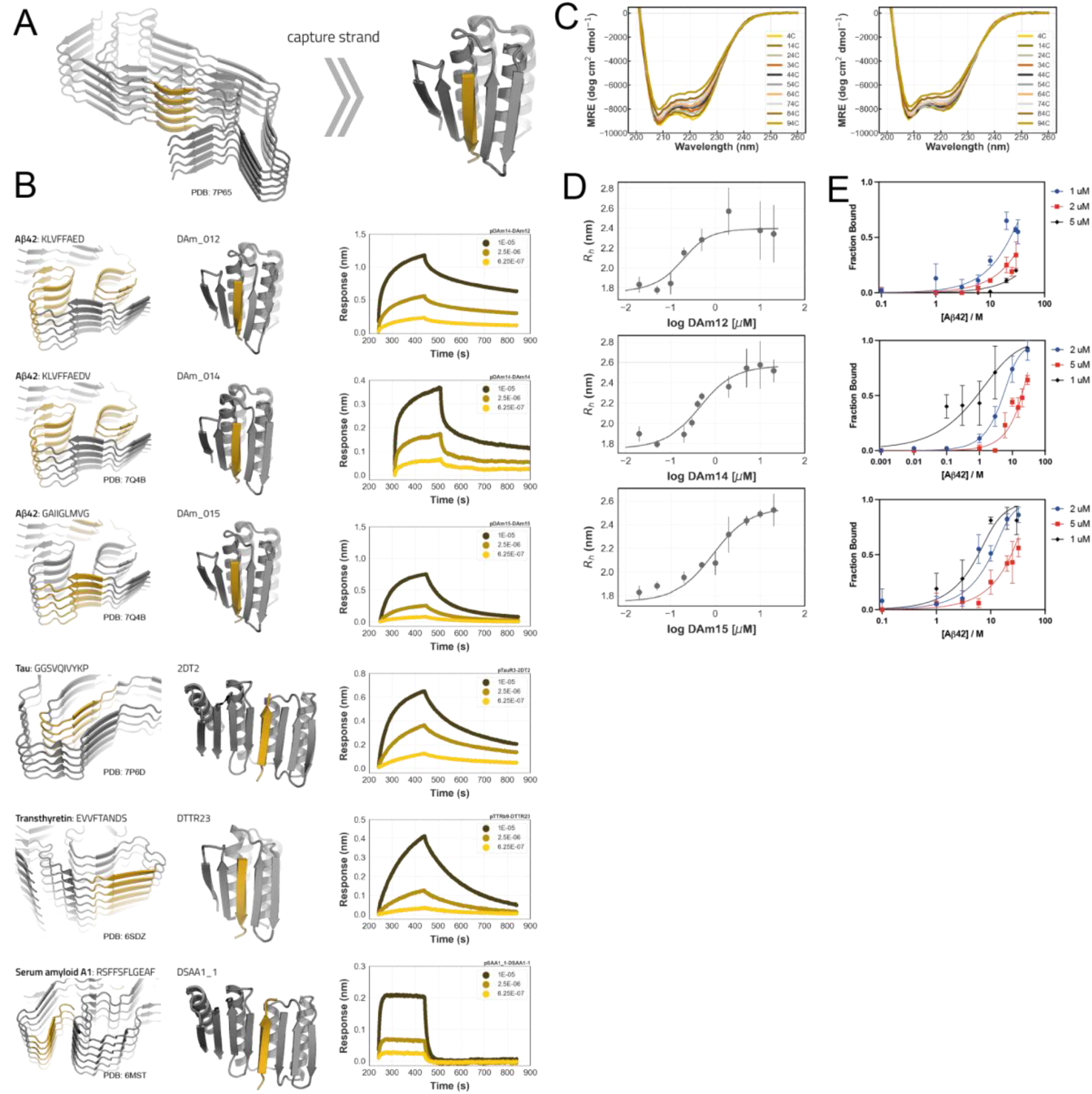
Design of amyloid peptide traps. **a:** To modulate fibril formation we design binders that can sequester a region that participates in fibril formation (yellow strand left Tau fibril). **b:** Models of designed proteins (middle) that bind amyloidogenic fragments (left yellow strand) of five different amyloid forming proteins in BLI experiments (right). **c:** Circular dichroism temperature melt spectra of DAm14 and DAm15. **d:** Microfluidic diffusional sizing (MDS) binding isotherms of DAm12, DAm14 and DAm15 binding to Aβ42 monomers. **e:** MDS binding of pre-formed Aβ42 fibrils to designs DAm11 (top), DAm14 (middle) and DAm15 (bottom).

To design such binders, we started from the design constraint that the peptide side chains facing the core of the binding scaffold must be primarily hydrophobic; since the peptide is bound in a β-strand conformation, every other residue is in the core and hence must be hydrophobic. We scanned the primary sequences of the Abeta peptide, Microtubule associated protein Tau, transthyretin and serum amyloid A1 for regions that matched this pattern (fig s10). Matched regions were docked in a β-conformation into the binding cleft of the scaffolds, and the scaffold interface residues were redesigned to maximize contacts to the amyloid derived β-strand, including surface-exposed hydrophobic interactions as described above. Designs with docked peptides predicted to participate in fibril or oligomer formation based on experimentally determined amyloid structures (Gremer et al. 2017; Shi et al. 2021; Y. X. Jiang et al. 2022; Guerrero-Ferreira et al. 2018; Schmidt et al. 2019; Liberta et al. 2019) were selected for experimental characterization.

The amyloid strand binders were first tested using the bicistronic expression screen described above; amyloid peptide fragments were fused to the C-terminus of GFP and co-expressed with polyhistidine tagged binder. After IMAC purification, we found using SDS PAGE that peptides derived from Aβ42, Transthyretin, Tau and Serum amyloid A1 interacted with the binders. In SEC, binder and peptide fusion protein co-eluted indicating the complexes remain stably associated even when diluted on the column. The designed scaffolds were also stable and mostly monodisperse by SEC when purified in absence of their target peptides (figure s11). We synthesized biotinylated versions of the single strand Aβ42, Transthyretin, Tau and Serum amyloid A1 fragments targeted by the designs and immobilized them on streptavidin biosensors to test in BLI. All purified designs bound their target peptides (figure 4b); we also observed some cross-reactivity consistent with similarities in the amyloid forming sequences. For example, the Aβ42 binders DAm14 and DAm15 bind their target and also interact with peptides derived from Transthyretin and Tau (Fig s11). Circular dichroism spectroscopy and SEC experiments indicated DAm14 and DAm15 were folded and thermostable, indicating that the promiscuous binding was not due to protein unfolding (figure 4c). Other designs such as the transthyretin binder DTTR23, Tau binder 2DT2 and serum amyloid A binder DSAA1_1 were more selective (figure s11 and s12) towards their targets.

We next investigated the binding properties of DAm12, DAm14 and DAm15 to the Aβ42 monomer using Microfluidic diffusional sizing (MDS). Measurements indicated that DAm12 and DAm14 interacted with the monomeric form of the Aβ42 peptide with dissociation constants of 83 and 350 nM whereas the Kd for DAm_015 was 755 nM (figure 4d and table s4). The designs could also interact with pre-formed Aβ42 fibrils (figure 4e).

After characterizing the binding interaction between the binders and their targets we next investigated the effect of the designs on amyloid fibril formation. To this end we tested Aβ42 fibril formation in the presence of DAm_012, DAm_014 and DAm_015 in a Thioflavin T (ThT) assay. We observed robust fibril formation in the control reactions (Fig 5a-c and fig s14) but in presence of the designs fibril formation was significantly retarded in a concentration dependent manner, with DAm_012 and DAm_014 being more potent than DAm_015 consistent with the tighter dissociation constants measured through MDS (figure 5a-c and figure 4d and table s4). DAm14 and DAm15, at stoichiometric ratios, completely inhibited fibril growth for at least 30h. DAm_012 prevented detectable amyloid formation for 10h even under a 1:2 sub-stoichiometric ratio of inhibitor to peptide, comparable to clinical stage therapeutic antibodies raised against this same target, including one approved drug, aducanumab (fig 5d) (Linse et al. 2020). Like DAm14 and DAm15, DAm12 is thermostable and remains folded up to 94°C in CD spectroscopy (figure s11 and s13). In a control experiment, C104 (figure 2a) and a previously *de novo* designed binder with a mixed α/β topology (Sahtoe et al. 2021) showed significantly lower inhibitory potential, indicating that the presence of a hydrophobic cleft surrounded by β-sheet structure is insufficient for inhibition (fig s14b-d).

**Figure 5.**
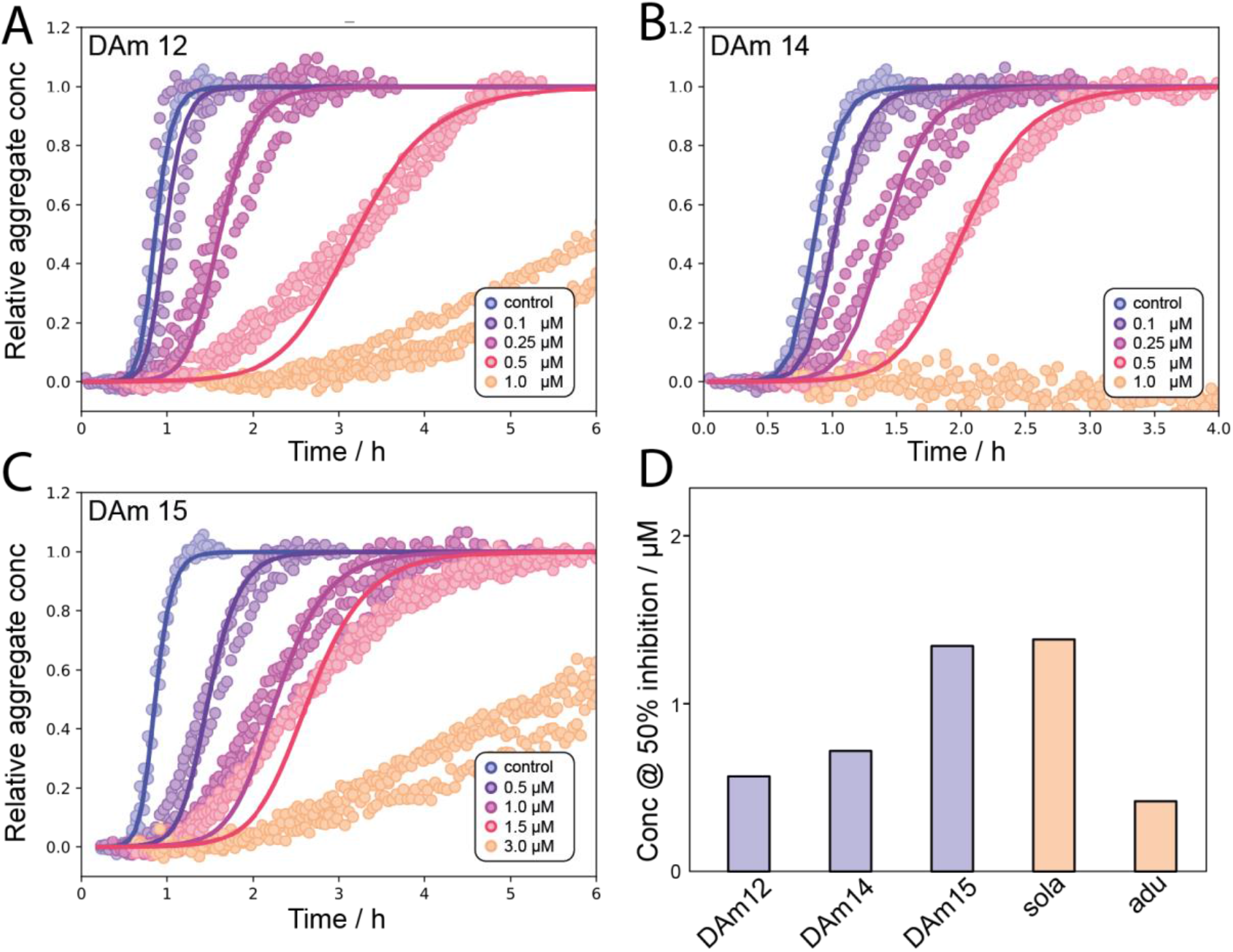
Inhibition of fibril formation. **a-c:** Aβ42 binders DAm12, DAm14 and DAm15 strongly inhibit fibril formation at sub-micro molar concentrations in a ThT aggregation assay. **d:** The inhibitory potential of the designed binders and clinical antibodies against Aβ42 aggregation is compared by evaluating the concentration of inhibitor at which the aggregation reaction has been slowed by a fixed amount (i.e. the half time of aggregation is increased by 50%). Lower values indicate higher potency. The values for the clinical antibodies solanezumab (sola) and aducanumab (adu) are obtained from (Linse et al. 2020). Points are ThT fluorescence measurements, solid lines are fits of the kinetics expected when inhibitor binds monomer with the above measured affinity and also inhibits aggregation by direct interactions with the aggregates.

In order to understand in more detail the mechanistic drivers of the observed inhibition, we used kinetic modeling (Meisl et al. 2016) to dissect the overall changes in the aggregation profiles in terms of changes in the molecular rate constants. Chemical kinetic analysis shows that there is a contribution to the aggregation behavior from the direct binding to the monomeric precursor peptide. The analysis further shows that there is an additional contribution to the retardation of the aggregation process from inhibition of fibril-related molecular steps. Indeed, the binders are more potent than would be expected based on their monomer binding ability alone. Even if all monomer sequestered by the binders is unable to take part in the aggregation reaction, the resulting slow down of aggregation is significantly less than observed in experiment. Only when we additionally allowed the compound to interact with the aggregated species, directly slowing the rate of aggregation, were we able to fully account for the observed inhibitory effect (see fitted curves in fig 5).

The presence of an inhibitory mechanism that includes this type of interaction with aggregated species is also supported by the observation that the binders also can interact directly with fibrils by MDS (fig 4e).

## Discussion

We present a general approach for designing binders targeted to disordered stretches of proteins and peptides that can adopt β-strand or β-hairpin conformations. The designed binders are folded and bind the target peptides with nanomolar affinities *in vitro* and in cells and can be incorporated into larger assemblies through fusion of peptide or binder to other components. Binding hydrophobic regions of proteins is challenging because the properties that make proteins stick to hydrophobic surfaces can also lead to poor solubility and highly indiscriminate binding; the overall geometry of the designed binding pocket and the dynamic sheet opening/closure observed by NMR appear to limit such adverse effects. While the apo state is dynamic, the x-ray crystal structure of a designed binder-peptide complex is highly ordered and very close to the design model.

The highly specific shape complementary binding pockets in our designs nearly completely engulf the bound peptide. This enables capture of protein segments that are prone to amyloid formation such as those found in Amyloid precursor protein, Microtubule associated protein Tau, transthyretin and serum amyloid A1. The designs potently inhibit the formation of the Aβ42 fibrils that are a hallmark of Alzheimer’s disease (AD), at a similar potency as clinically evaluated antibodies, including an approved drug (aducanamab). This result is particularly significant since it is challenging to elicit antibodies to monomeric forms of peptides which spontaneously self associate, and structurally our β-sheet clamping approach can likely generate more extensive interactions with extended β-strand peptides than antibody loops. The designs may also be useful in blocking smaller amyloidogenic oligomers, as the oligomers use similar stretches of sequences to self-assemble and are considered highly toxic precursors to fibrils. Moving forward, the designs should be useful for testing hypotheses on the pathogenesis of AD and other amyloid diseases, and could also contribute to new diagnostic and therapeutic approaches.

## Materials and methods

### Protein design

#### Backbone generation

We explored two approaches to generate scaffolds with β-sheets with open slots for peptide β-strand insertion (figure 1b and s1) using blueprint based backbone building in (Py)Rosetta (Koga et al. 2012; Lin et al. 2015; Huang et al. 2011; Chaudhury, Lyskov, and Gray 2010; Leman et al. 2020). In a first two-domain binder approach (figure s1a), we started from a scaffold, 2003285_0000, designed by Fold-It players (Koepnick et al. 2019) (domain 1) and generated a β-sheet that extends from the C-terminal strand of the scaffold using blueprint based backbone generation (Huang et al. 2011; Koga et al. 2012). In the next step this sheet was further expanded into a second mixed alpha/beta domain with three strands and one helix or four strands and two helices. The central strand of the β-sheet that encompasses both domains was split off from generating an individual peptide in β-strand conformation that can bind the designed deep cleft between domain 1 and domain 2. A connecting loop linking the helices that make up the interdomain interface was next generated using loop closure (Brunette et al. 2015) to yield a single polypeptide two-domain binder that clamps the peptide on either side through β-strand backbone hbonds (figure s1a). The same approach was followed to generate β-hairpin binding scaffolds.

In the second approach, a different foldit scaffold, 2003333_0006 (Koepnick et al. 2019), was modified to function as a peptide binder (figure s1b). The connection between β-strand 3 and 4 was removed to create the individual peptide component. To stabilize the modified binder and ensure its solubility in absence of the peptide, we designed buttressing secondary structure elements that support the binding interface and scaffold. β-strand 3 was paired with another antiparallel strand whereas helices 1 and 2 were backed up by either one or two supporting helices. After backbone generation, Rosetta combinatorial sequence design calculations were used to optimize the sequences of both the scaffold and the peptide for high affinity binding. Designs with favorable interaction energy, few unsatisfied buried polar atoms and high shape complementarity, and for which Rosetta folding simulations yielded models close to the designed model were selected for experimental characterization.

#### Sequence design

The amino acid sequence of the newly built polyvaline backbones were optimized using Rosetta flexible backbone enabled combinatorial side chain design followed by a second design round for the peptide-binder interface (Bhardwaj et al. 2016; Hosseinzadeh et al. 2017). Ref2015, beta_nov16 or beta_genpot scorefunctions were used during design (Alford et al. 2017). For a subset of designs, buried polar hydrogen bond networks were designed using the HBNet mover (Boyken et al. 2016).

The affinity between peptide and binder was computationally improved by introducing hydrophobic interaction pairs to the solvent exposed side of the interface. All solvent exposed interactions pairs for which the Cα atoms were within 6 Å from each other were selected and allowed to be redesigned with the PackRotamersMover to only Phe, Ala, Met, Ile, Leu, Tyr, Val and Trp using a fixed backbone. For the computational affinity optimization of the natural target peptides, all surface exposed residues on only the binder within 6 Å of the target hydrophobic side chain were allowed to be redesigned. Residues around the redesigned interactions pairs were repacked. Single redesigned pairs and combinations of pairs were selected for experimental characterization.

In order to facilitate crystallization, the surface residues outside the interface were redesigned using ProteinMPNN (Dauparas et al. 2022) for design C104. The structure of sequences obtained from ProteinMPNN were predicted using AlphaFold2 (Jumper et al. 2021) and designs with rmsd <= 1.5 and plDDT >= 85 to the original designed model were selected for experimental characterization.

#### Design of rigid helical fusions

Rigid fusions of peptide binders and components of the LHD hetero-oligomer system was performed as described previously (Hsia et al. 2021; Sahtoe et al. 2022).

#### Matching natural peptide sequences to scaffolds

The protein sequences of Amyloid precursor protein, Microtubule associated protein Tau, Transthyretin and Serum amyloid A1 were searched for burial patterns that are also present in the peptides of designs C34, C37, C104 and CH15. For C104 both the designed model and the crystal structure of C104, minimized with FastRelax (Tyka, Jung, and Baker 2012), was used. The burial patterns representing relative positions of solvent inaccessible residues versus solvent accessible residues in the designed peptides were identified by visual inspection. For each peptide, all amyloidogenic protein sequence-frames of length n, where n is the number of residues in the designed peptide, were scanned for matching regions.

Only residues Phe, Ala, Met, Ile, Leu, Val or Gly were allowed at the solvent inaccessible positions. At the remaining positions, all residues were allowed except for Pro which was only allowed at either terminus. When a match was identified, the sequence of the template designed peptide was mutated to the sequence of the matched sequence of the amyloidogenic protein. The resulting peptide-binder complex was minimized and the residues in the interface of the designed binder were redesigned to optimally match the amyloidogenic sequence by also including hydrophobic interaction pairs across the solvent accessible area of the interface (see above).

PDB models of the designed proteins and example scripts can be downloaded as supplementary files.

### Protein expression and purification

Synthetic genes encoding designed proteins were purchased from Genscript or Integrated DNA technologies (IDT) in the pET29b expression vector or as eBlocks (IDT) and cloned into customized expression vectors (Wicky et al. 2022) using golden gate cloning. A His6x tag was included either at the N-terminus or at the C-terminus as part of the expression vector. In some cases a TEV protease recognition site was introduced at the N-terminus after the histidine tag. Peptide genes were purchased as fusion proteins to either the C-terminus of sfGFP or the N-terminus of a ubiquitin-AviTag-His6x construct separated by a Pro-Ala-Ser linker. Bicistronic genes were ordered as described (Sahtoe et al. 2022). Detailed construct information is provided in the supplementary information.

Proteins were expressed using autoinducing media consisting of TBII media (Mpbio) supplemented with 50×5052, 20 mM MgSO_4_ and trace metal mix in BL21 LEMO E.coli cells. Proteins were expressed under antibiotic selection at 37 degrees Celsius overnight or at 18-25 degrees Celsius overnight after initial growth for 6-8h at 37 degrees Celsius. Cells were harvested by centrifugation at 4000x g and resuspended in lysis buffer (100 mM Tris pH 8.0, 200 mM NaCl, 50 mM Imidazole pH 8.0) containing protease inhibitors (Thermo Scientific) and Bovine pancreas DNaseI (Sigma-Aldrich) before lysis by sonication. One millimolar of the reducing agent TCEP was included in the lysis buffer for designs with free cysteines.

Proteins were purified by Immobilized Metal Affinity Chromatography. Cleared lysates were incubated with 2-4ml nickel NTA beads (Qiagen) for 20-40 minutes before washing beads with 5-10 column volumes of lysis buffer, 5-10 column volumes of high salt buffer (10 mM Tris pH 8.0, 1 M NaCl) and 5-10 column volumes of lysis buffer. Proteins were eluted with 10 ml of elution buffer (20 mM Tris pH 8.0, 100 mM NaCl, 500 mM Imidazole pH 8.0). His6x tags were cleaved by dialyzing IMAC elutions against 20 mM Tris pH 8.0, 100 mM NaCl, 1 mM TCEP overnight in the presence of His6x tagged TEV protease followed by a second IMAC column to remove His6x-TEV and uncleaved protein.

Single cysteine variants of DAm12, DAm14 and DAm15 where purified as described above and labeled with Alexa488-C5-maleimide (Thermo) at a concentration of between 50-100 μM of protein and a 2-5 fold molar excess of label in SEC buffer supplemented with 1 mM TCEP protected from light. After 3h at room temperature or overnight at 4 degrees Celsius the labeling reaction was quenched by the addition of 1M DTT.

All protein preparations were as a final step polished using size exclusion chromatography (SEC) on either Superdex 200 Increase 10/300GL or Superdex 75 Increase 10/300GL columns (Cytiva) using 20 mM Tris pH 8.0, 100 mM NaCl. The reducing agent TCEP was included (1 mM final concentration) for designs with free cysteines. For designs where a substantial void volume peak was present in addition to the monomer peak, the monomer peak was pooled and reinjected. Only designs where upon reinjection the void peak was mostly absent were further pursued. SDS-PAGE and LC/MS were used to verify peak fractions. Proteins were concentrated to concentrations between 0.5-10 mg/ml and stored at room temperature or flash frozen in liquid nitrogen for storage at -80. Thawing of flash frozen aliquots was done at room temperature or 37 degrees Celsius. All purification steps from IMAC were performed at ambient room temperature.

The C104.1 complex was prepared by incubating binder with a 3-5 fold molar excess of the peptide for 3h at room temperature followed by SEC.

### Peptide synthesis

All Fmoc-protected amino acids were purchased from P3 Bio. The biotinylated peptides obtained by synthesis were padded at the C-terminus with SGGSGGKbiotin where Kbiotin is a Fmoc-Lys(Biotin)-OH building block also purchased from P3 Bio. Oxyma was purchased from CEM; DIC from Oakwood Chemicals. DMF was purchased from Fisher Scientific and treated with an Aldraamine trapping pack (Sigma-Aldrich) prior to use. Piperidine was purchased from Sigma-Aldrich. Cl-TCP(Cl) resins were purchased from CEM. The peptides were synthesized on a 0.1mmol scale using microwave-assisted solid-phase peptide synthesis via a CEM LibertyBlue system, then subsequently cleaved with a cleavage cocktail consisting of TFA, TIPS, water, and DODT (92.5:2.5:2.5:2.5 in order). The cleavage solution was concentrated *in vacuo*, precipitated into cold ether, and spun down by way of centrifugation. This pellet was washed and spun down again with ether (2x), then dried under nitrogen, resuspended in water and ACN, and purified by RP-HPLC on an Agilent 1260 Infinity Semi-prep system with a gradient from 20% to 70% over a period of 15min (A: H2O with 0.1% TFA, B: ACN with 0.1% TFA). The purified peptide fractions were combined into one, lyophilized, and massed in a tared scintillation vial for the final product. Peptides derived from Transthyretin, Tau, and Serum amyloid A1 were purchased from WuXi. Depending on the isoelectric point, lyophilized peptides were solubilized in buffers containing either 100 mM Tris pH 8.0 or 100 mM MES pH 6.5 and stored at -20 degrees Celsius.

### Mammalian cell culture and transfection

HeLa cells (ATCC CCL-2) were cultured in Dulbecco’s modified Eagle’s medium (DMEM) (Gibco) supplemented with 1 mM L-glutamine (Gibco), 4.5 g/liter D-glucose (Gibco), 10% fetal bovine serum (FBS), and (1×) nonessential amino acids (Gibco). Cells were kept in culture at 37°C and 5% CO2 and split twice per week by trypsinization using 0.05% trypsin EDTA (Gibco) followed by passage at 1:5 or 1:10 into a new tissue culture (TC)–treated T75 flask (Thermo Scientific ref 156499). Before transfection, cells were plated at 20,000 cells per well in Cellview cell culture slides (Greiner Bio-One ref 543079) for 24 hours after which transfection took place using 187.5 ng total DNA per well and 1 μg/μl PEI-MAX (Polyscience) mixed with Opti-MEM medium (Gibco). Transfected cells were incubated at 37°C and 5% CO2 for 24 to 36 hours before being imaged.

### Fluorescent microscopy

Three dimensional images were acquired with a commercial OMX-SR system (GE Healthcare) using a 488 nm Toptica diode laser for excitation. Emission was collected on a PCO.edge sCMOS cameras using an Olympus 60× 1.42NA PlanApochromat oil immersion lens. 1024×1024 images (pixel size 6.5 μm) were captured without binning. AcquireSR Acquisition control software was used for data collection. Z-stacks were collected with a step size of 500 nm and 15 slices per image. The images were deconvolved with an enhanced ratio using SoftWoRx 7.0.0 (GE Healthcare). Finally, cell images were sum projected using Fiji v2.1.0. Scale bars equal 10 microns.

### Biolayer interferometry

Biolayer interferometry experiments were performed on an OctetRED96 BLI system (ForteBio, Menlo Park, CA) at room temperature in Octet buffer (10 mM HEPES pH 7.4, 150 mM NaCl, 3 mM EDTA, 0.05% surfactant P20) supplemented with 1mg/ml bovine serum albumin (SigmaAldrich). Prior to measurements, streptavidin-coated biosensors were first equilibrated for at least 10 min in Octet buffer. Chemically synthesized peptides with a C-terminal biotin or enzymatically biotinylated peptide-fusion proteins (see supplementary spreadsheet for details) were immobilized onto the biosensors by dipping them into a solution with 100 to 500 nM protein until the response reached between 10% and 50% of the maximum value followed by dipping sensors into fresh octet buffer to establish a baseline for 60 s. Titration experiments were performed at 25 degrees Celsius while rotating at 1000 rpm. Association of designs was allowed by dipping biosensors in solutions containing designed protein diluted in octet buffer until equilibrium was approached followed by dissociation by dipping the biosensors into fresh buffer solution to monitor the dissociation kinetics. In the peptide binding cross specificity assays each biotinylated peptide was loaded onto streptavidin biosensors in equal amounts followed by 2 min of baseline equilibration.

Then association and dissociation with all the different binders was allowed for 400 s for each step. For the designed peptide-binder pairs, binder concentrations were around the Kd of the interaction between the loaded peptide and its designed binding partner whereas the concentrations for the amyloid binders were 10, 2.5 and 0.625 μM. Global kinetic or steady-state fits were performed on buffer subtracted data using the manufacturer’s software (Data Analysis 9.1) assuming a 1:1 binding model.

### Enzymatic biotinylation of proteins

Proteins with Avi-tags (GLNDIFEAQKIEWHE; see supplementary materials) were purified as described above and biotinylated *in vitro* using the BirA500 (Avidity, LLC) biotinylation kit. 840 ul of protein from an IMAC elution was biotinylated in a 1200 μl (final volume) reaction according to the manufacturer’s instructions. Biotinylation reactions were allowed to proceed at either 4°C overnight or for 2-3 hours at room temperature on a rotating platform. Biotinylated proteins were purified using SEC on a Superdex 200 10/300 Increase GL (GE Healthcare) or S75 10/300 Increase GL (GE Healthcare) using SEC buffer (20 mM Tris pH 8.0, 100 mM NaCl).

### Circular Dichroism Spectroscopy

CD spectra were recorded in a 1 mm path length cuvette at a protein concentration between 0.3-0.5 mg/mL on a J-1500 instrument (Jasco). For temperature melts, data were recorded at 222 nm between 4 and 94 °C every 2 C°, and wavelength scans between 190 and 260 nm at 10 C° intervals starting from 4 C°. Experiments were performed in 20 mM Tris pH8.0, 20 mM NaCl. The high tension (HT) voltage was monitored according to the manufacturers recommendation to ensure optimal signal-to-noise ratio for the wavelengths of interest.

### SEC binding assays

SEC binding assays between purified designs and GFP-peptide fusions were performed on a Superdex 75 increase 10/300 GL (Cytiva) in 20 mM Tris pH 8.0, 100 mM NaCl using 500 ul injections containing 15 or 20 μM final concentration of each component. Binding reactions were allowed to equilibrate for at least 45 minutes before injection. For the subunit exchange experiment, the disulfide stabilized complex between C104.2 and ubiquitin-pep104.2 as well as the control base non-covalent complex were allowed to form overnight at a 20 μM equimolar concentration under oxidizing conditions after which competing GFP-pep104 was added to the pre-formed complexes to a final concentration of 20 μM. After at least 45 minutes the reaction was injected on SEC. Elution profiles were collected by monitoring absorbance at 230 nm and 395 nm (absorbance of GFP). All experiments were performed at room temperature.

### Disulfide formation assay

Individual protein components were purified as described above in the presence of 1 mM TCEP except for in the last SEC step where no reducing agent was present. Reactions were incubated at room temperature using 50 μM of each component in 20 mM Tris pH 8.0, 100 mM NaCl. Reactions were stopped by adding an equal volume of 2x non-reducing SDS protein loading buffer at the indicated time points.

### NMR

All NMR experiments for C34 were performed on Bruker Avance III HD 14.1 T or 18.8 T spectrometers equipped with cryogenically cooled, x,y,z pulse-field gradient triple-resonance probes. Resonance assignments were obtained by triple-resonance (HB)CBCA(CO)NNH, HNCACB, HNCO, HN(CA)CO, and HNN experiments (Sattler, Schleucher, and Griesinger 1999) acquired using U-{^13^C,^15^N}-labeled samples. Note that the spectra shown in figure 3A were recorded at 25°C, but resonance assignment for free C34 was done at 50°C to reduce the line broadening arising from conformational exchange. ^15^N *R*_2_ rates for the bound state of C34 were measured using the in-phase CPMG experiment (Hansen, Vallurupalli, and Kay 2008) with v_CPMG_ = 1 kHz, T_relax_ = 30 ms and CPMG refocusing pulses applied at a γB_1_/2π = 5.7 kHz field and phase-modulated according to the {x,x,y,-y} cycling scheme (B. Jiang et al. 2015). A NOESY dataset for recording intermolecular NOEs was acquired as previously described (Zwahlen et al. 1997) with a mixing time of 150 ms, using 450 μM of U-{^13^C,^15^N}-labeled C34 and 450 μM of unlabeled peptide at natural isotopic abundance.

### Amyloid-β expression, purification and labeling

Amyloid-β (Aβ1-42) peptide was expressed and purified as reported previously (Abelein et al. 2020). In short, the synthetic gene coding for NT*FlSp was purchased from GenScript (GenScript Biotech, Netherlands), ligated into pT7 plasmid containing TEV recognition site (TRS) for Aβ42 (Kronqvist et al. 2017), and transformed into chemically competent E. coli BL21 (DE3) cells and expressed as described earlier (G. Chen et al. 2017). Upon cleavage of the fusion protein with TEV protease, the sample was dissolved in 15 mL 8 M guanidine-hydrochloride (GuHCl) and monomeric Aβ purified on a Superdex 30 26/600PG size exclusion column, and lyophilized as aliquots until further use. To generate fibrils, several aliquots of lyophilised Aβ were combined for an increased protein concentration by dissolving in 1 ml of 8 GuHCl, and subjected to SEC on a Superdex 75 10/300 Increase column in 20 mM sodium phosphate, 0.2 mM EDTA buffer at pH 8.0. Subsequently collected monomeric peptide, typically at a concentration of 30 μM, was pipetted into PEGylated plates (Corning 3881) and incubated at 37°C in a plate reader, with 100 rpm orbital shaking.

To track the degree of monomer conversion into fibrils, ThT was added exclusively to control wells, and the fibrils were harvested from ThT-free sample wells after the plateau was reached in the control wells.To perform binding experiments of monomeric Aβ with the binders, cysteine-carrying Aβ mutant (S8C) has been expressed and purified as described previously (Thacker et al. 2022). Briefly, the plasmid carrying synthetic genes with E. coli optimized codons for S8C mutant (developed by Thacker and colleagues and purchased from Genscript) were transformed into BL21 DE3 pLysS star E. coli strain and the protein was expressed in auto-induction medium (Studier 2005). Upon purification using IEX and subsequent SEC on a 26 × 600 mm Superdex 75 column, S8C monomer was eluted in a sodium phosphate buffer supplemented with 3 mM DTT to prevent its dimerization, and lyophilized. For conjugation of the protein with a fluorescent dye, the lyophilized fractions were dissolved in 8 M GdnHCl and subjected to SEC in buffer without DTT, before adding the Alexa Fluor 488 dye (Thermofisher) in 5x< molar excess. The protein-dye mixture was incubated overnight at 4 °C, the free dye removed via column chromatography, and the protein used immediately.

### Kinetic assays of fibril inhibition

Aliquots of purified lyophilized Aβ were dissolved in 8 M GuHCl, and the monomeric protein was isolated by gel filtration on a Superdex 75 10/300 Increase column in 20 mM sodium phosphate, 0.2 mM EDTA buffer at pH 8.0. Samples were prepared on ice, using careful pipetting to avoid introduction of air bubbles, and pipetted into a 96-well half-area plate of PEGylated black polystyrene with a clear bottom (Corning 3881), 100 μl per well, three to four replicates per sample. All samples diluted with buffer to the final concentration of 2 μM Aβ were supplemented with 6 μM ThT (Sigma), with a range of concentrations of the binders per experiment. The kinetic assays were initiated by placing the 96-well plate at 37 °C under quiescent conditions in a plate reader (FLUOstar Optima BMGLabtech). The ThT fluorescence was measured through the bottom of the plate every 165s s with a 440 nm excitation filter and a 480 nm emission filter.

### Analysis of aggregation kinetics

Integrated rate laws describing the aggregation of Aβ42 were derived previously (Cohen et al. 2013). They reproduce well the kinetic curves obtained in ThT assays and can be used to quantify inhibitory effects. Here, we used the amylofit platform (Meisl et al. 2016) to determine the rate constants of aggregation in the absence of inhibitor. Using the affinities of binder to monomer determined by MDS, we then calculated the concentrations of monomer expected to be bound at each binder concentration. Assuming all monomer bound is completely removed from the aggregation reaction (i.e. ignoring dissociation of the monomer-binder complex over the timescale of aggregation), the effect of binders on the aggregation reaction is the same as a lowering of the monomer concentration. The kinetic curves resulting from this effective reduction of monomer concentration were then computed using the amylofit platform (see figure s14), and the effect was found to be insufficient to explain the observed degree of inhibition. We then explored if the presence of an additional mechanism of inhibition, by interaction with aggregated species, was able to describe the observed aggregation. To model this additional inhibition we allowed the rate of secondary nucleation to vary with binder concentration, as detailed previously (Meisl et al. 2016). These results are shown as solid lines in Fig. 5 and can describe well the inhibition at substoichiometric binder concentrations. At higher binder concentrations, when the majority of monomer is expected to be bound, these fits perform less well and thus only the experimental measurements, not the fits, are shown at the highest binder concentrations.

### Microfluidic diffusional sizing

Binding affinity of the binders and monomeric Aβ was measured on a Fluidity One-M (Fluidic Analytics). Fluorescently labeled Aβ mutant was mixed with unlabeled binders at a range of concentrations and incubated on ice for at least 30 min. Before the measurements, microfluidic circuits of the Fluidity-One M chip plate were primed using sample buffer. To create a binding curve for individual designs, each one of the different Aβ-binder mixtures was measured in triplicates. KD values were determined by non-linear least squares fitting as described previously (Schneider et al. 2021) using Prism (GraphPad Software). For microfluidic diffusional sizing experiments concerning interactions of binders with Aβ fibrils, microfluidic devices have been fabricated and operated as described previously (Arosio et al. 2016; Qin, Xia, and Whitesides 2010). In brief, the microfluidic devices were fabricated in PDMS by standard soft-lithography techniques and bonded onto a glass coverslip after activation with oxygen plasma. Sample loading from reservoirs connected to the respective inlets and control of flow rate was achieved by applying negative pressure at the outlet using a glass syringe (Hamilton) and a syringe pump (neMESYS, Cetoni GmbH). Images were recorded using a custom-built inverted epifluorescence microscope fitted out with a fluorescent filter set with an excitation filter at 475 ± 35 nm, emission filter at 525 ± 30 nm and dichroic mirror for 506 nm (Laser 2000) for detection of Alexa-488 labeled binders. Images were taken using Micro Manager, typically at flow rates 60 and 100 μL/h, and lateral diffusion profiles were recorded at four different positions along the microfluidic channels. Diffusion profiles extracted from fluorescence images and confocal recordings were fitted using a custom-written analysis software by numerical model simulations solving the diffusion– advection equations for mass transport under flow (Müller et al. 2015).

### Crystal structure determination

The C104.1 complex was (19 mg/ml) crystallized using the vapor diffusion method at room temperature in 0.1 M Tris pH 7.8, poly-γ-glutamic acid low molecular weight polymer, 15% PEG 4000 (Molecular dimensions) before the crystals were harvested in 25% glycerol as a cryoprotectant. Data was collected at the Advanced Photon Source at Argonne National Laboratory. Diffraction images were integrated using XDS (Kabsch 2010) or HKL3000 (Otwinowski and Minor 1997) and merged/scaled using Aimless (Winn et al. 2011). Starting phases were obtained by molecular replacement using Phaser (McCoy et al. 2007) using the computational design models of the individual N and C terminal domains of C104.1 as search models. Structures were refined using either phenix.refine (Adams et al. 2010) or Refmac (Murshudov, Vagin, and Dodson 1997) and PDB-REDO (Joosten et al. 2014).

Model building was performed using COOT (Emsley and Cowtan 2004). Lack of density at the C-terminus of the peptide prompted us to examine the possibility of a β-strand register shift for the peptide binding. OMIT maps were used to decrease the model bias. In addition, the peptide was modeled in several off-target β-strand registers. Overall refinement statistics and B-factors, were better for the model where the peptide was modeled in the designed on-target β-strand register. The final model was evaluated using MolProbity (Williams et al.

2018). Data collection and refinement statistics are recorded in Table 3. Data deposition, atomic coordinates, and structure factors reported in this paper have been deposited in the Protein Data Bank (PDB), http://www.rcsb.org/ with accession code 8FG6.

## Acknowledgements

We thank the Institute for Protein Design and Baker lab members for general discussion and in particular Yakov Kipnis, Inna Goreshnik, Wei Yang and Gyu-Rie Lee and Sam Pellock for helpful advice,Patrick Salveson and Lance Stewart for discussions on amyloidogenic peptides and neurodegenerative diseases, Nathan Ennist for assistance with circular dichroism spectroscopy, Basile I.M. Wicky and Lukas Milles for providing golden gate cloning vectors, Lauren Carter for assistance with international shipping. This work was supported by a gift from Gates Ventures (D.D.S., D.B.), the Audacious Project at the Institute for Protein Design (H.L.H, H.C., J.D., H.N.,D.B.), a gift from Amgen (M.A., D.B.), a grant from DARPA supporting the Harnessing Enzymatic Activity for Lifesaving Remedies (HEALR) Program (HR001120S0052 contract HR0011-21-2-0012, X.L., A.K.B., D.B.), Canadian Institutes of Health Research grant (FND-503573, L.E.K.) and Natural Sciences and Engineering Research Council of Canada grant (2015-04347 L.E.K.). Crystallographic data was collected at the Advanced Photon Source (APS) Northeastern Collaborative Access Team beamlines, which are funded by the National Institute of General Medical Sciences from the National Institutes of Health (P30 GM124165). This research used resources of the Advanced Photon Source, a U.S. Department of Energy (DOE) Office of Science User Facility operated for the DOE Office of Science by Argonne National Laboratory under Contract No. DE-AC02-06CH11357.

## Author contributions

D.D.S. designed research, developed computational design approach, performed design calculations and analyzed data. D.D.S. and H.L.H. characterized proteins. E.A.A., M.M.S. and G.M. performed fibril inhibition assays, microfluidic diffusional sizing and kinetic fitting of inhibition data under supervision of T.P.J.K. A.K.B., H.N., A.K. and D.D.S. determined crystal structure. E.R. performed NMR experiments and analyses under supervision of L.E.K. P.L., M.L. and X.L. synthesized peptides. M.A. and J.D. performed and analyzed mammalian cell based experiments. D.D.S. and D.B. wrote the manuscript with input from all authors. D.B. supervised research.

---

**Table S1.**
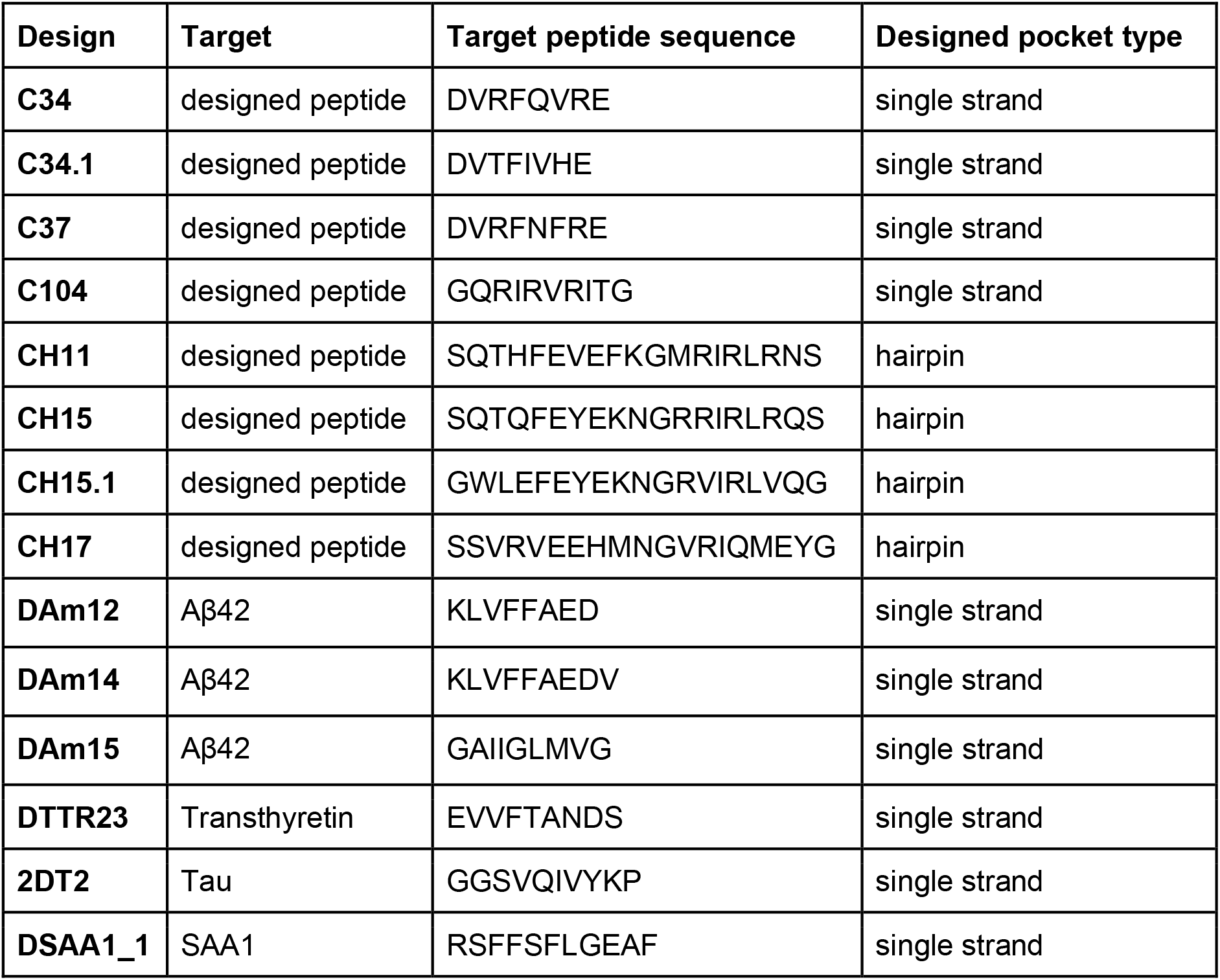
Overview designs.

**Table S2.** Biolayer interferometry global kinetic fitting parameters.

| | $K_D$ (μM) | $k_{on}$ ($M^{-1} s^{-1}$) | $k_{off}$ ($s^{-1}$) | chi-sqr | R-sqr |
| --- | --- | --- | --- | --- | --- |
| **C34** | $12.8 \pm 0.38$ | $1.37 \cdot 10^4 \pm 391$ | $0.176 \pm 0.0016$ | 0.15 | 0.99 |
| **C37** | $0.31 \pm 6.9 \cdot 10^{-4}$ | $547.4 \pm 1.1$ | $1.7 \cdot 10^{-4} \pm 1.8 \cdot 10^{-7}$ | 8.7 | 0.99 |
| **C104** | $0.157 \pm 0.0015$ | $6348 \pm 56$ | $9.96 \cdot 10^{-4} \pm 3.9 \cdot 10^{-6}$ | 0.48 | 0.99 |
| **CH11** | $0.167 \pm 2.78 \cdot 10^{-3}$ | $4990 \pm 59.1$ | $8.3 \cdot 10^{-4} \pm 9.7 \cdot 10^{-6}$ | 0.11 | 0.99 |
| **CH15** | $44 \pm 11.2$ | $434 \pm 10.9$ | $0.019 \pm 9.9 \cdot 10^{-5}$ | 0.02 | 0.99 |
| **CH17** | $9.4 \pm 0.31$ | $1155 \pm 39$ | $0.0146 \pm 3.11 \cdot 10^{-4}$ | 0.064 | 0.99 |
| **C34.1** | $2.3 \pm 0.017$ | $1358 \pm 9.9$ | $3.08 \cdot 10^{-3} \pm 6.38 \cdot 10^{-6}$ | 7.34 | 0.99 |
| **CH15.1** | $0.05 \pm 4.1 \cdot 10^{-4}$ | $6730 \pm 34.4$ | $3.3 \cdot 10^{-4} \pm 2.2 \cdot 10^{-6}$ | 0.46 | 0.99 |

**Table S3.** Crystallographic data collection and refinement.

|  |  |
| --- | --- |
|  | **C104.1 (PDB code: 8FG6)** |
| **Data Collection** |  |
| Space group | P 31 2 1 |
| *Cell dimensions* |  |
| *a, b, c* (Å) | 64.54, 64.54, 112.61 |
| $\alpha, \beta, \gamma$ (°) | 90, 90, 120 |
| Resolution (Å) | 55.89 - 2.30 (2.53 - 2.30) |
| $R_{\text{merge}}$ | 0.118 (0.966) |
| $R_{\text{pim}}$ | 0.0338 (0.2644) |
| $I/\sigma(I)$ | 12.97 (2.87) |
| $CC_{1/2}$ | 0.996 (0.984) |
| Completeness (%) | 99.50 (99.51) |
| Redundancy | 13.6 (14.3) |
| **Refinement** |  |
| Resolution (Å) | 55.89 - 2.30 (2.53 - 2.30) |
| No. reflections | 12514 (3052) |
| $R_{\text{work}} / R_{\text{free}}$ | 0.2552 (0.3551)/ 0.2917 (0.3573) |
| *No. atoms* |  |
| Protein | 1288 |
| Water | 13 |
| Ramachandran<br>Favored/allowed<br>Outlier (%) | 96.79/ 3.21/ 0.00 |
| *R.m.s. deviations* |  |
| Bond lengths (Å) | 0.013 |
| Bond angles (°) | 1.65 |
| $B_{\text{factors}}$ (Å<sup>2</sup>) | |
| Protein | 50.91 |
| Water | 53.79 |

**Table S4.** Binding constants microfluidic diffusional sizing.

| Design | K<sub>D</sub> (nM) | 95%CI |
| --- | --- | --- |
| DAm11 | 3290 | 1240 - 8240 |
| DAm12 | 83 | 15.72 - 217.3 |
| DAm14 | 350 | 183.4 - 635.4 |
| DAm15 | 754 | 378.4 - 1386 |

**Fig S1.**
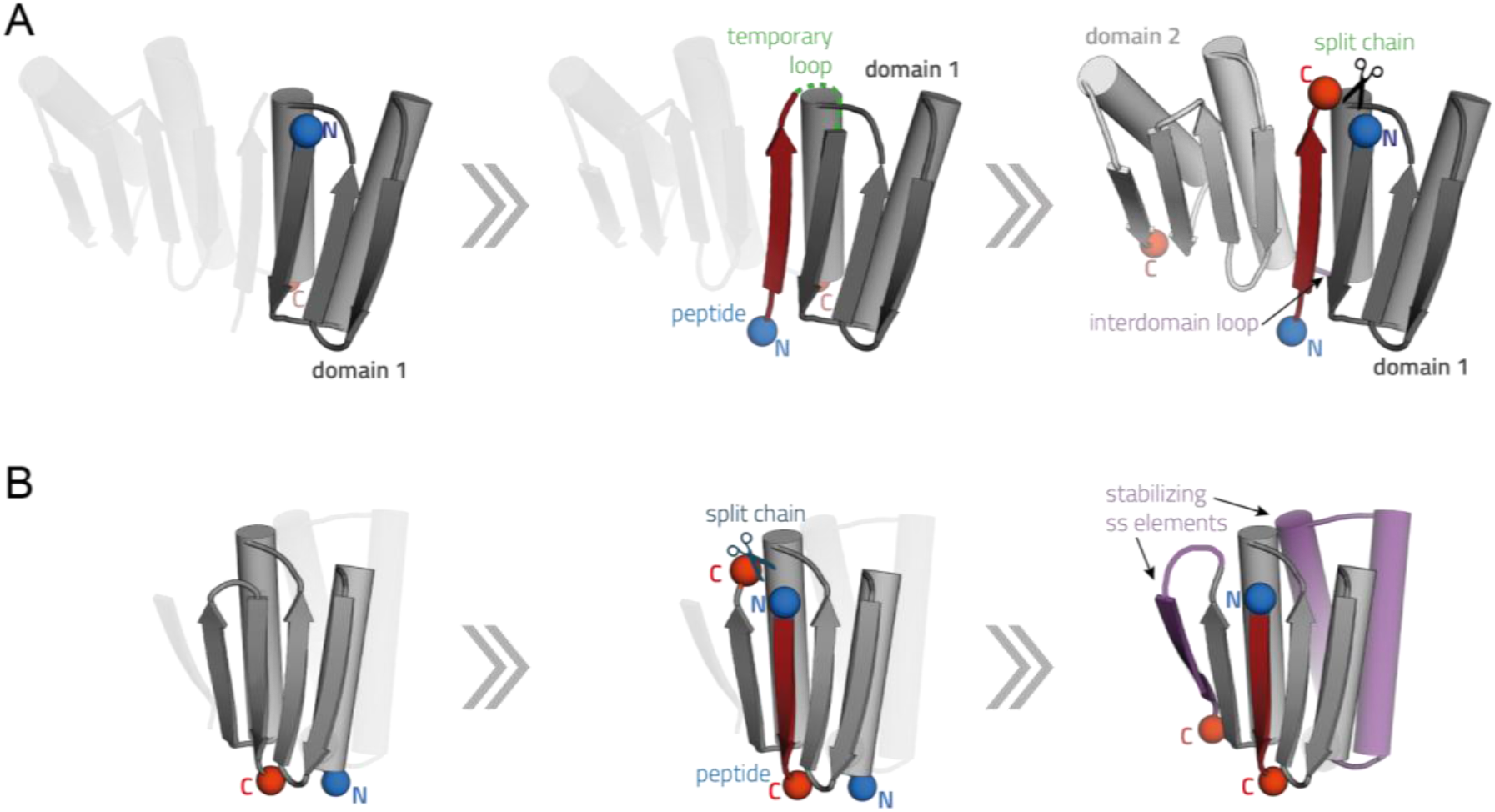
Design approach beta peptide binders. **a:** In approach 1 previously published (Koepnick et al. 2019) scaffold 2003285_0000 (gray, domain 1) was extended with a strand (dark red) and a globular alpha/beta domain (light gray) using blueprint based backbone building. The peptide was generated by introducing chain breaks. Loop closure between the C-terminus of domain 1 and N-terminus of domain 2 yields a single chain beta peptide binder in which the peptide complements the large beta sheet encompassing domains 1 and 2. **b:** In approach 2 a chain break was introduced in the connecting loop between strand 3 and 4 of the previously published scaffold 2003333_0006 (Koepnick et al. 2019) (gray) to generate the peptide (dark red). Stabilizing secondary structure elements (purple) were built using blueprint based backbone building at the N and C terminus of the scaffold.

**Fig S2.**
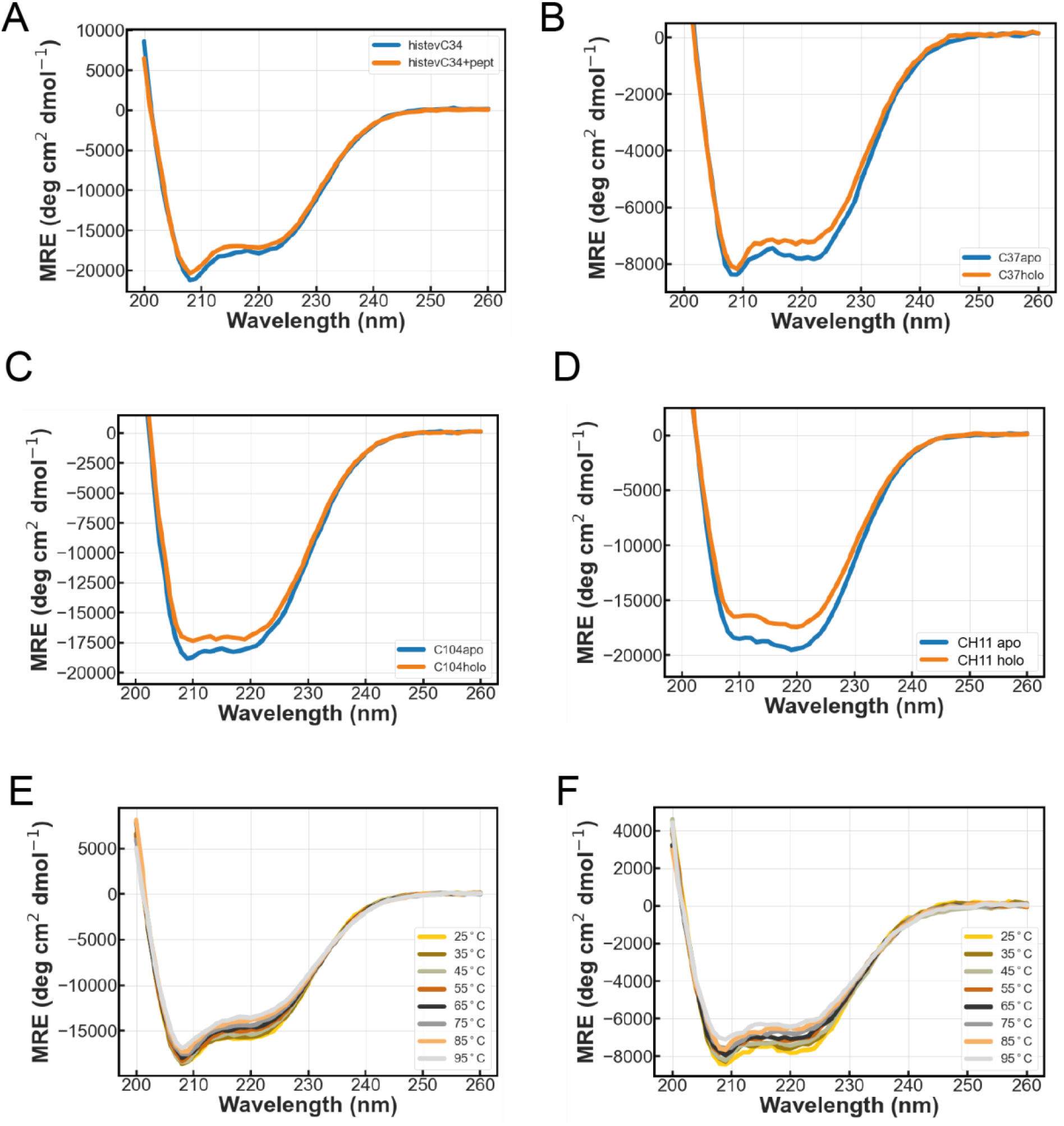
Biophysical characterization designs. **a-d:** Circular dichroism (CD) spectrum at 25**°**C of various binders with (blue) and without peptide (orange). **e:** CD spectra of C34 at different temperatures. **f:** CD spectra of C37 at different temperatures.

**Fig S3.**
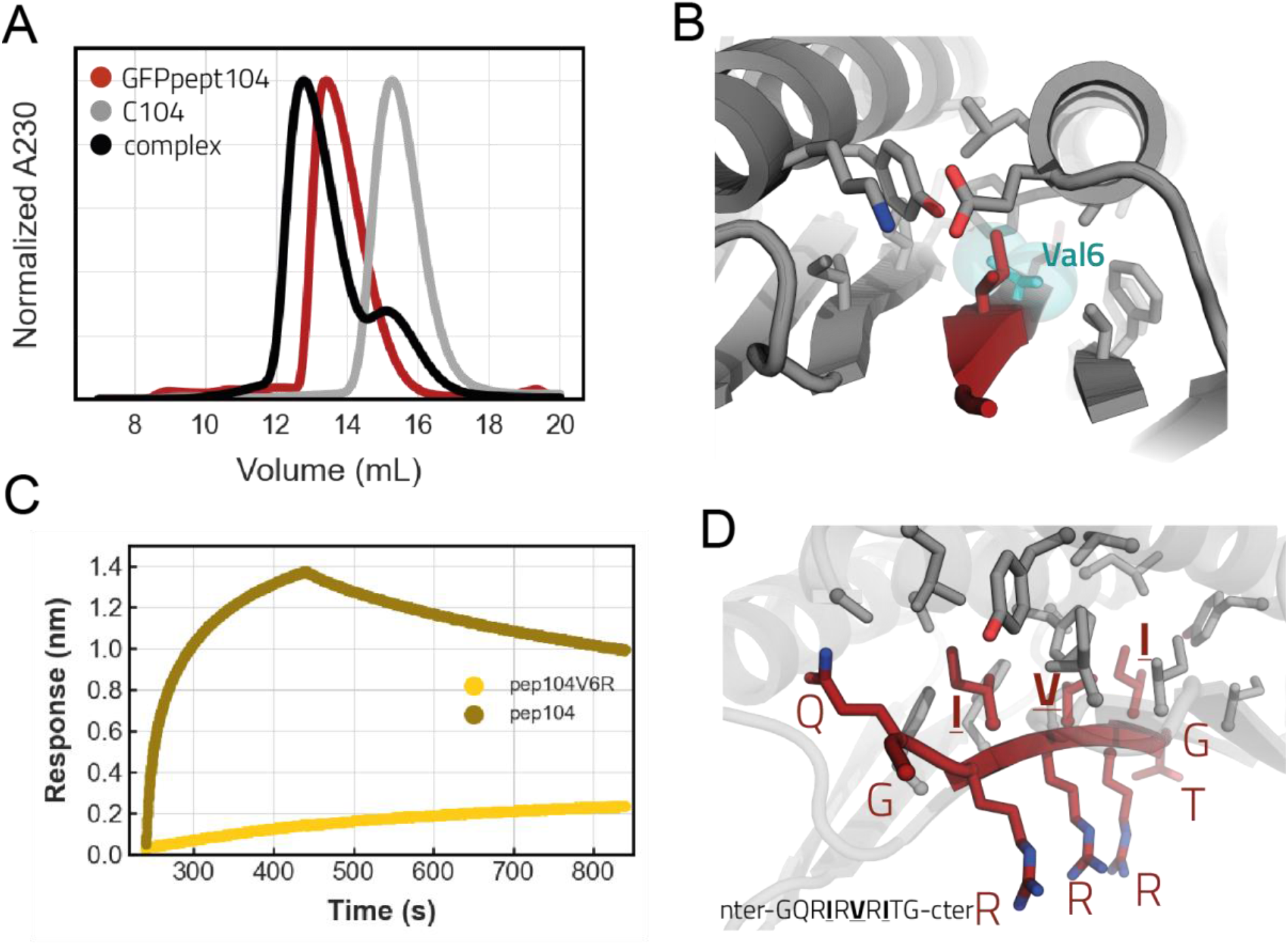
C104 controls. **a:** SEC binding assay showing that a fusion protein between GFP and 104 peptide binds to the C104 design on a S75 increase 10/300. **b:** Close-up view of the buried part of the C104 interface with Val6 shown in cyan sticks and spheres. Binder in gray and peptide in dark red. **c:** Biolayer interferometry trace of C104 binding to base peptide 104 and to a peptide with a V6R substitution. **d:** Interface close up view of C104 highlighting the hydrophobic-hydrophilic pattern of the peptide. Buried residues single letter amino acid identifiers are underlined.

**Fig S4.**
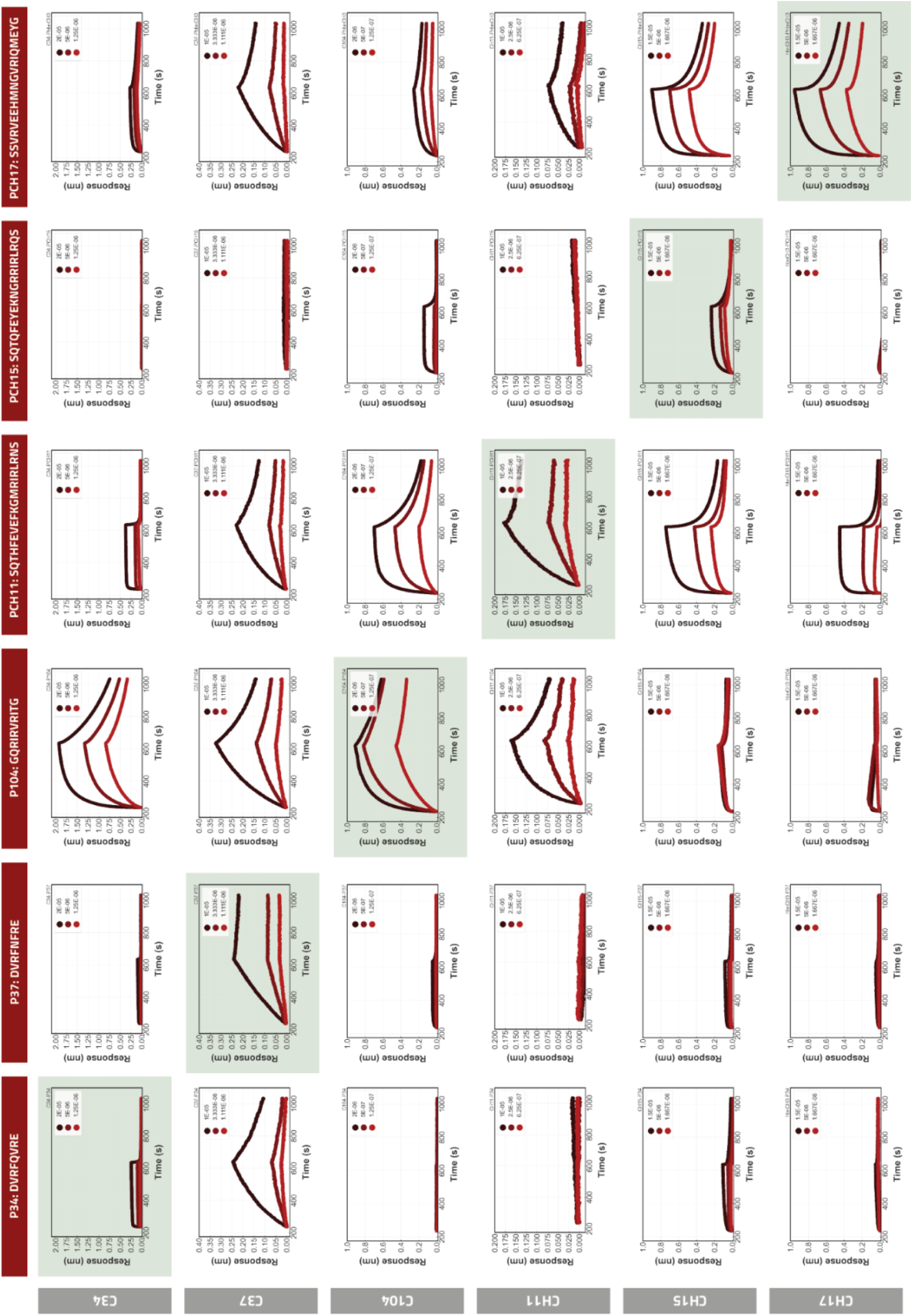
Specificity profile of peptide binder designs in BLI. Peptides were immobilized onto octet biosensors at equal densities and incubated with all designs in separate experiments at three different binder concentrations. The on-target interactions are indicated with a light green background. The experiment was done for each different peptide from the base designs (fig2a).

**Fig S5.**
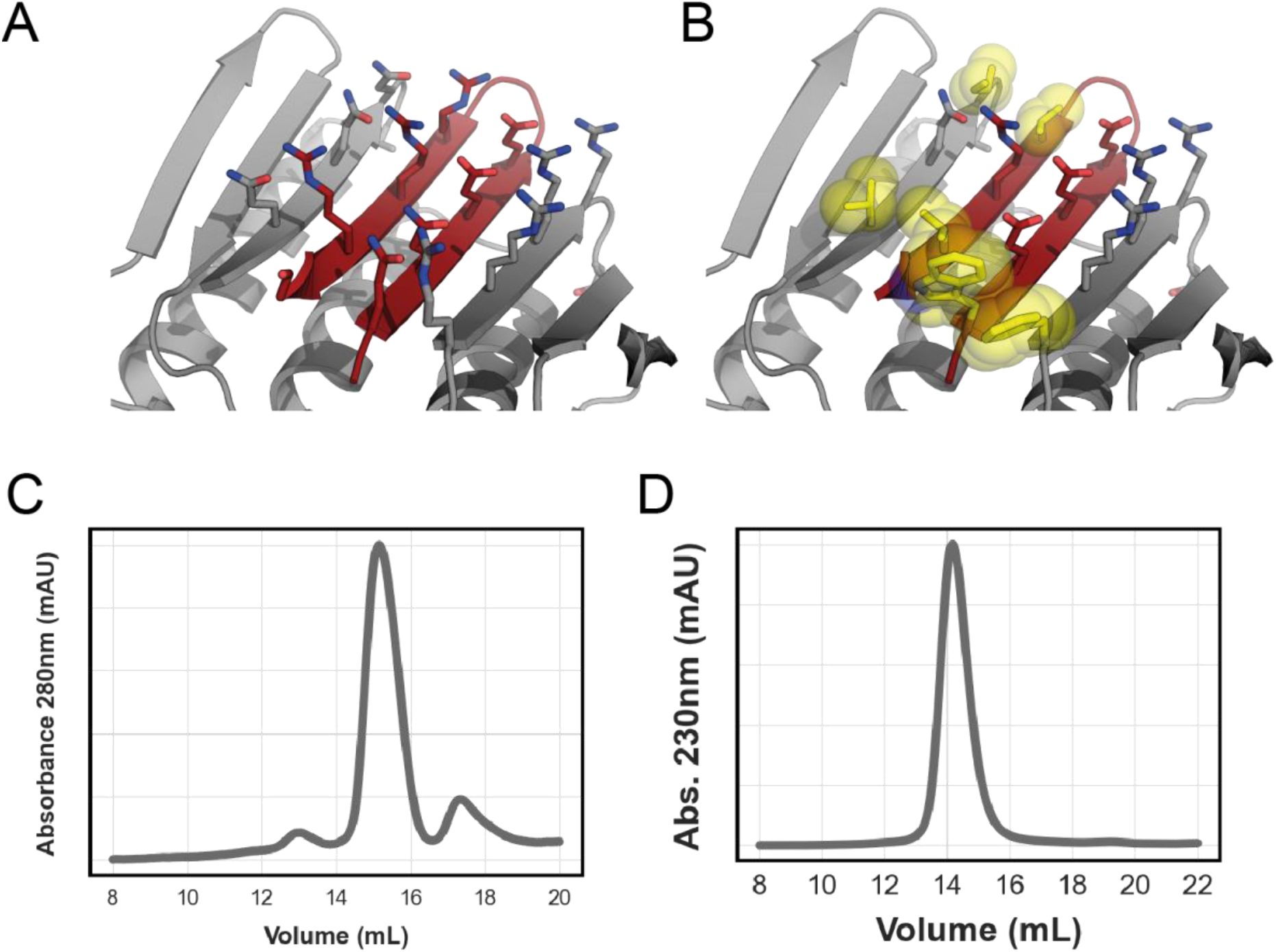
Computational affinity maturation by introducing solvent exposed hydrophobic interaction pairs. **a:** View of the solvent exposed interface of CH15 (binder gray, peptide dark red). **b:** View of the redesigned CH15.1 interface. Hydrophobic interaction pairs introduced to the base CH15 scaffold to improve affinity are highlighted in yellow sticks and spheres. Superdex 75 Increase 10/300 GL SEC traces of purified C34.1 **c)** and CH15.1 **d)**.

**Fig S6.**
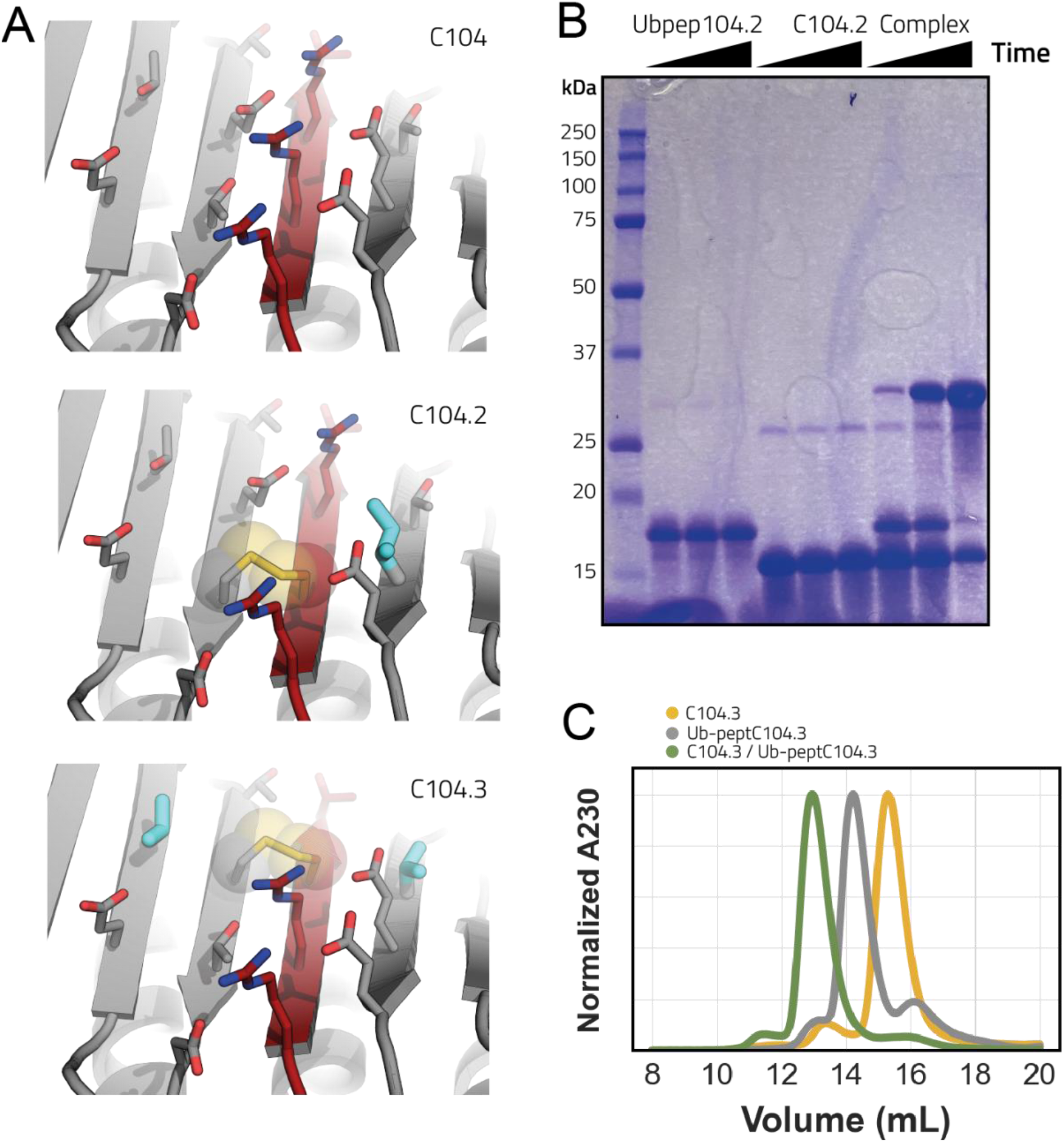
Disulfide functionalization of C104. **a:** Close-up of C104 surface exposed interface (top) and of the disulfide bridge variants C104.2 (middle) and C104.3 (bottom). Disulfide bonds are highlighted with spheres while additional redesigned residues to optimally accommodate the disulfide bridges are highlighted in cyan thicker sticks. Designed binder in gray and peptide in dark red. **b:** Coommassie stained non-reducing SDS-PAGE gel monitoring disulfide bridge formation of C104.2. Time points are t=0, t=90min and t=overnight. **c:** Superdex 75 increase 10/300 GL SEC binding assay confirming that the cysteine containing peptide of C104.3 fused to ubiquitin can bind to its designed cysteine containing binding partner C104.3.

**Fig S7.**
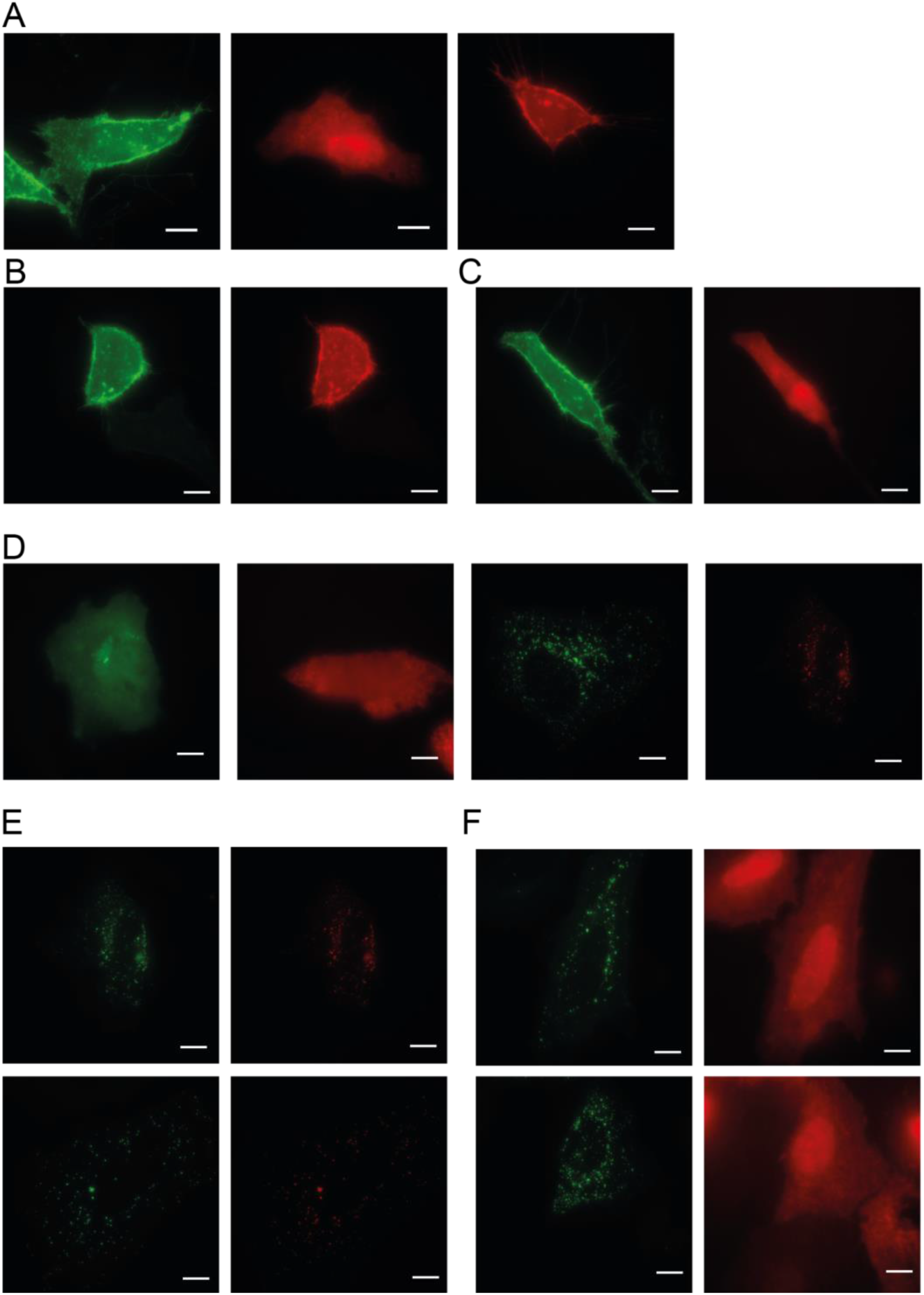
Fluorescent microscopy peptide-binder localization CH15.1. in HeLa cells. **a:** Full views fig 2j. **b:** Alternative view membrane localization. **c:** When F5K/L16K double mutant intended to disrupt binding is introduced to the peptide of CH15.1, CH15.1 binder fused to mScarlet (red channel) does not localize to the membrane anymore. **d:** Full views fig 2k. **e:** Alternative views of localization of CH15.1 mScarlet fusion to the two component GFP puncta. **f:** Two views showing that when the F5K/L16K double mutant is introduced to the peptide of CH15.1 the binder does not localize to the puncta anymore (red channels) even though the puncta still form (green channels). Scale bars 10 μm.

**Fig S8.**
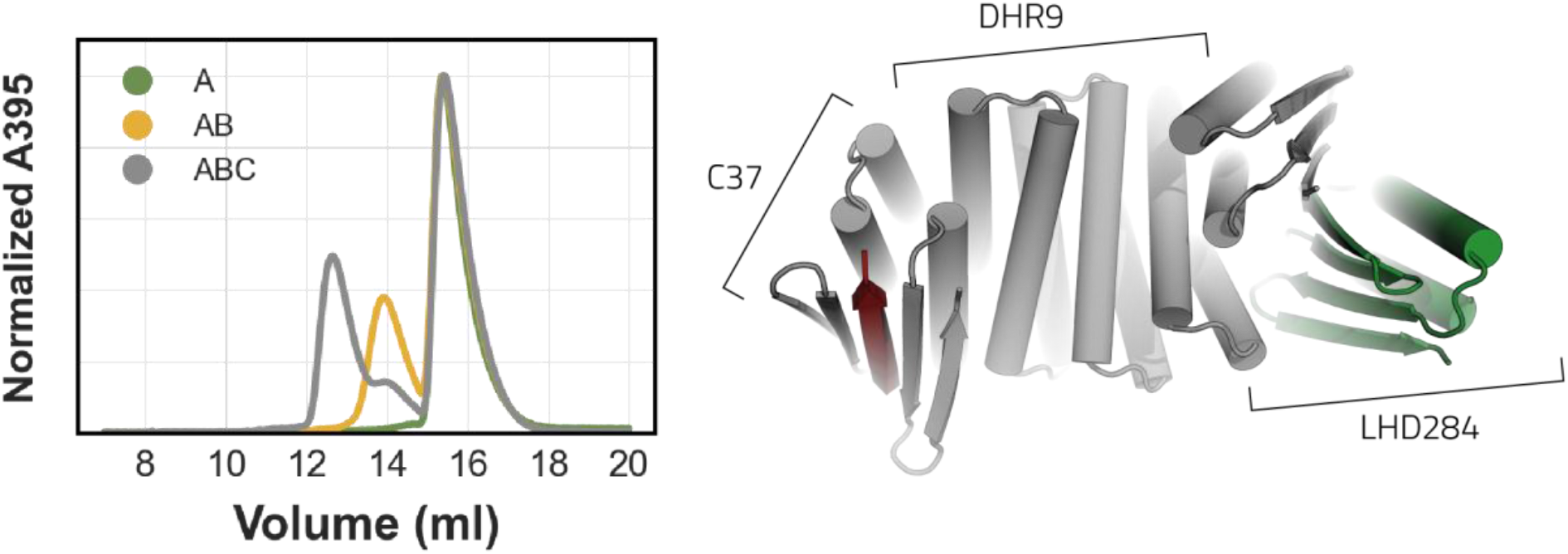
Incorporation of C37 into LHD hetero-oligomer system. Design C37 was rigidly fused to LHD284B_DHR9 (right) creating a single chain protein with two interfaces capable of binding the peptide of C37 and the designed binding partner of LHD284B_DHR9, LHD284A_DHR82. We validated the assembly of this ternary complex in a SEC binding assay on a S200 increase 10/300 GL. A: GFP-peptC37, B: GFP-peptC37 + LHD284B_DHR9, C: GFP-peptC37 + LHD284B_DHR9 + LHD284A_DHR82. Absorbance at 395 nm of the GFP-peptC37 was monitored to assess binding.

**Fig S9.**
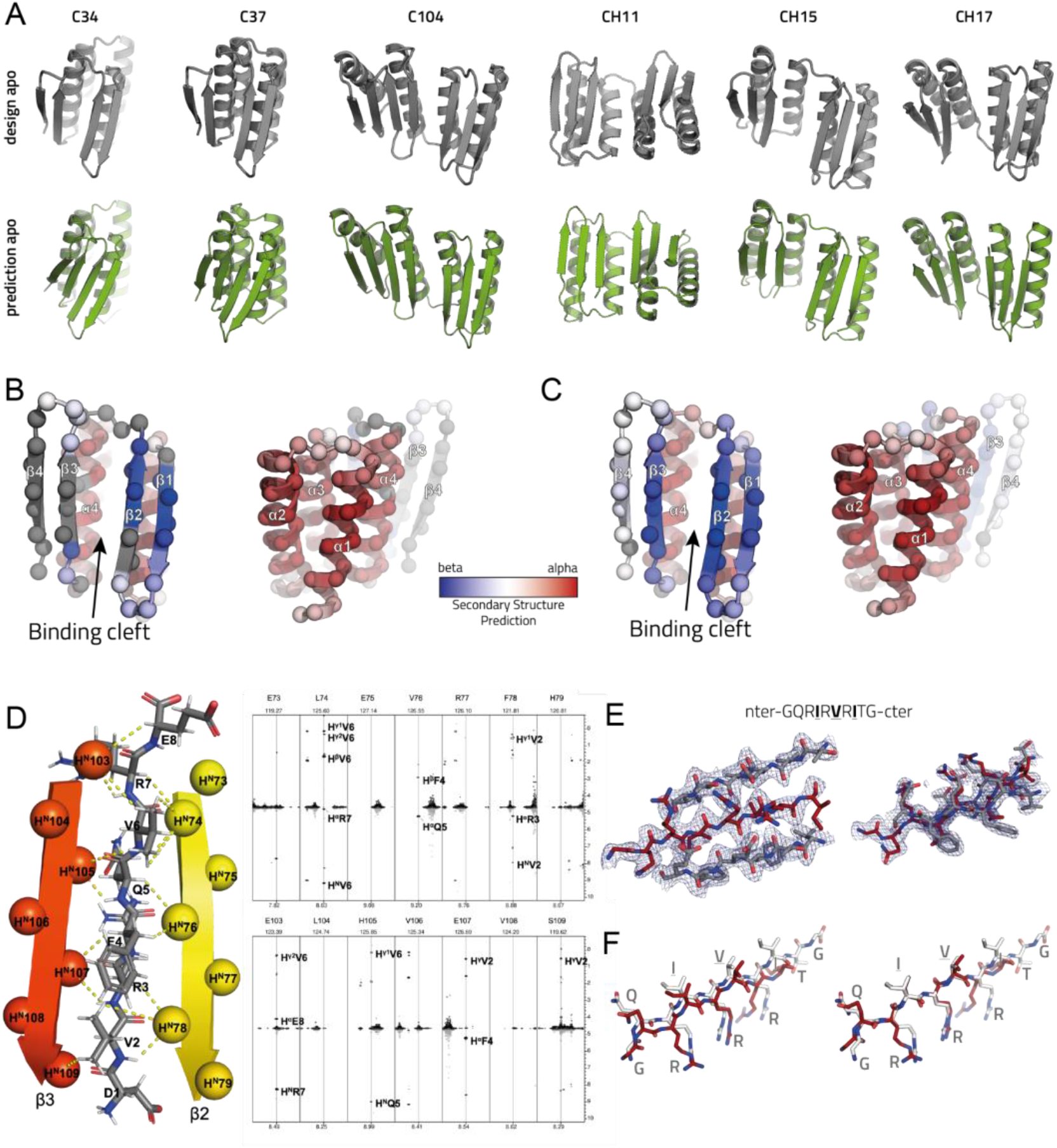
Structural characterization peptide binders. **a:** AlphaFold2 predictions of designed binder sequences in absence of peptide (bottom row) indicate closure of the binding pocket for some designs. **b:** Secondary structure prediction from NMR experiments on C34 apo mapped onto the cartoon model (two views) of C34 with C-alpha atoms shown as spheres (peptide not shown). No information available for residues in gray. These residues had broadened resonances due to conformational exchange. **c:** Same as **b)** but for C34 holo (peptide not shown). **d:** Intermolecular NOE contacts between C34 and the peptide measured as previously described (Zwahlen et al. 1997) using a sample comprised of a mixture of 450 μM U-{^13^C,^15^N} C34 and 450 μM unlabeled peptide. Strips from the 3D dataset are illustrated at the ^15^N chemical shifts of the amides of the indicated residue (top of panels) showing the detected intermolecular contacts between the amide protons of strands β2/β3 from C34 and the peptide (right panel). The protons linked via the observed NOEs are highlighted on the structure of the designed binder on the left panel. **e:** Two atomic views with 2mF_obs_−DF_calc_ electron density maps contoured at 1.5σ of the strand-strand interaction between the binder and peptide of C104. **f:** Left, View of peptide in designed model after superposition of entire designed (white) and xray structures. Right superposition on only peptides in designed and xray structure.

**Fig S10.**
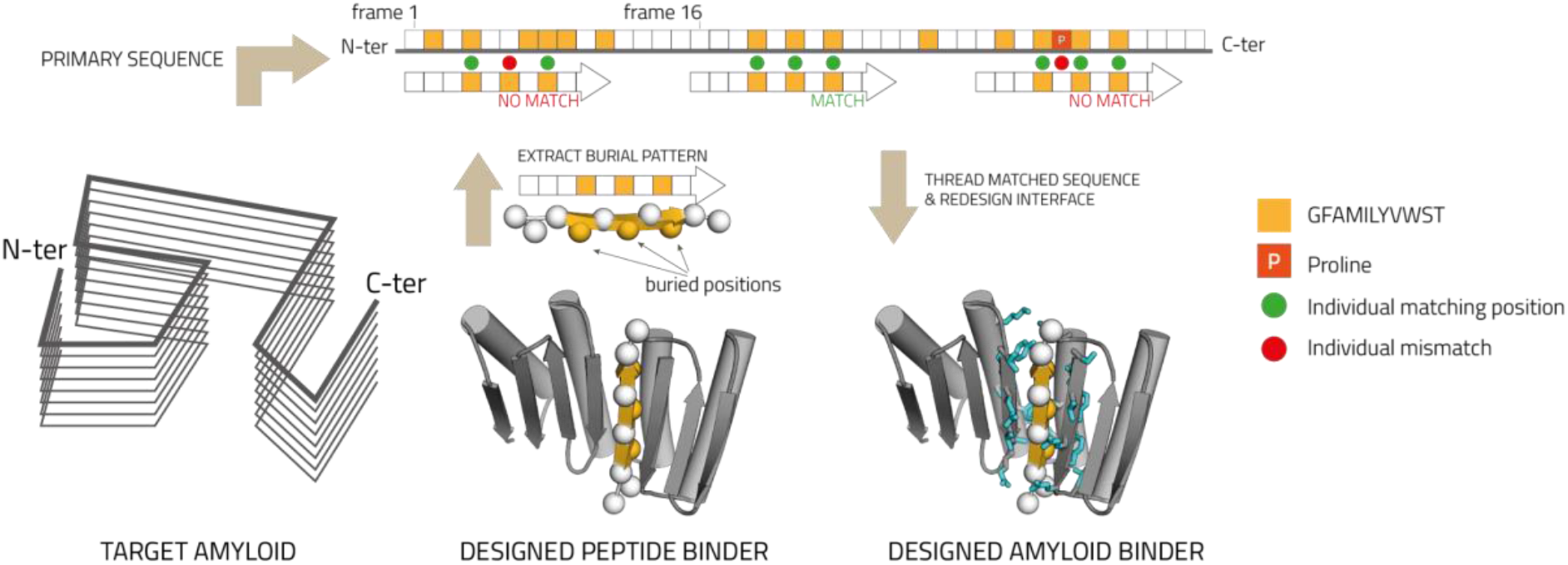
Amyloidogenic sequence docking. The designed peptides are optimized to only harbor GFAMILYVWST residues at the buried positions (yellow); charged residues cannot be accommodated at buried positions due to the high chance of burying a polar residue that cannot be satisfied by complementary side chains on the scaffold. To identify stretches of sequence present in amyloidogenic proteins that can be accommodated in a beta strand conformation in the binding pockets of the designs (fig 2a), the burial pattern of the peptides (middle) are matched to the primary sequence of the amyloidogenic protein (top). Because prolines disrupt beta conformation they are only allowed at termini. When a match is found the original peptide sequence is mutated to the matched sequence from the amyloid protein (threading) and docked back into the scaffold binding pocket followed by redesign of the scaffold interface residues (cyan sticks) to optimize interactions to the amyloidogenic sequence (right).

**Figure S11.**
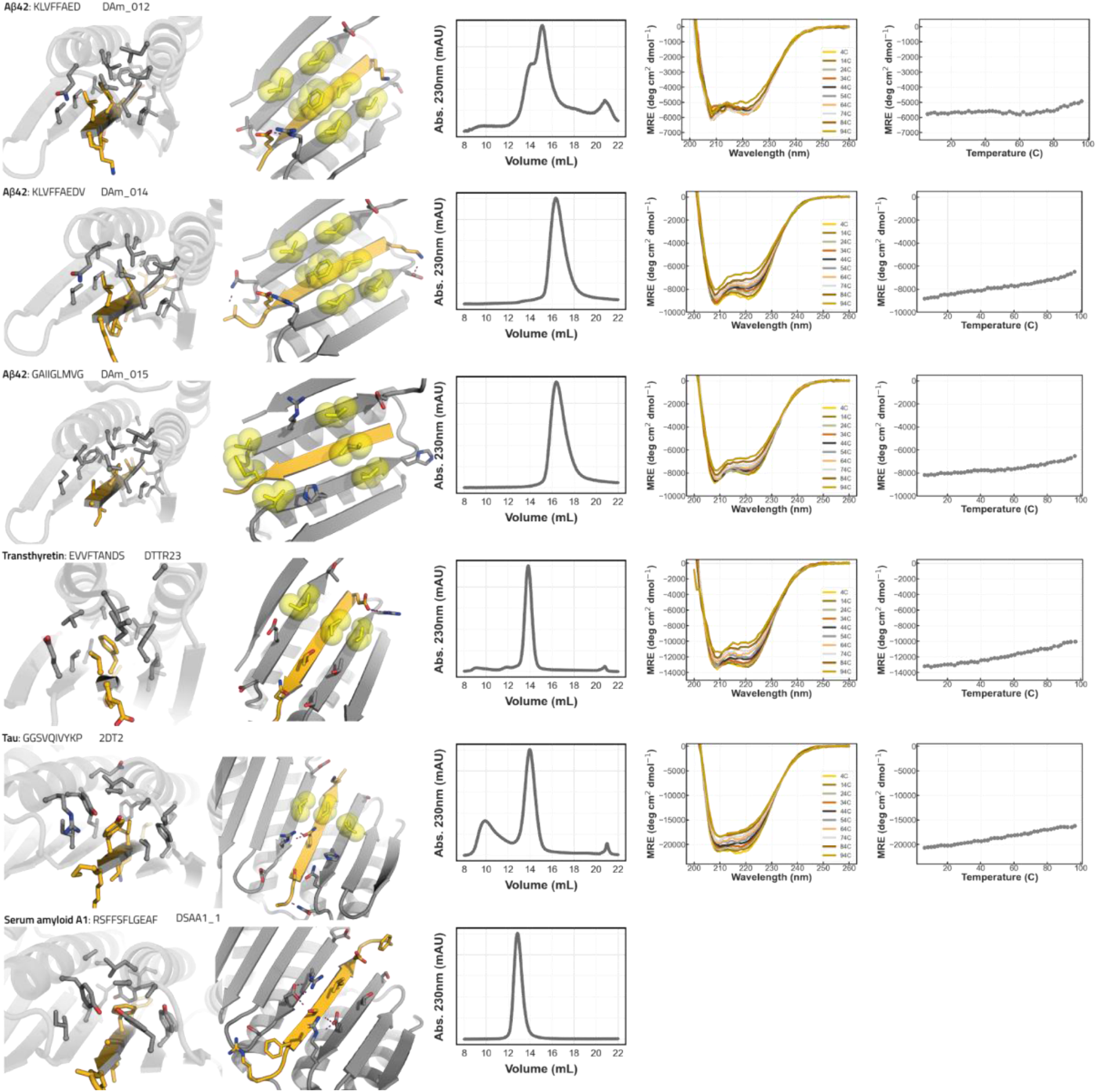
Characterization amyloidogenic strand binders. Close-up view of solvent inaccessible part interface (first column), close up view of solvent accessible part of interface with hydrophobic interaction pairs in yellow spheres and sticks (2nd column), SEC trace of binder on S75 increase 10/300GL (3rd column), CD wavelength scans (4th column) and CD temperature melt at 222 nm. CD wavelength scans for DAm14 and DAm15 are the same as in main fig 4c.

**Fig S12.**
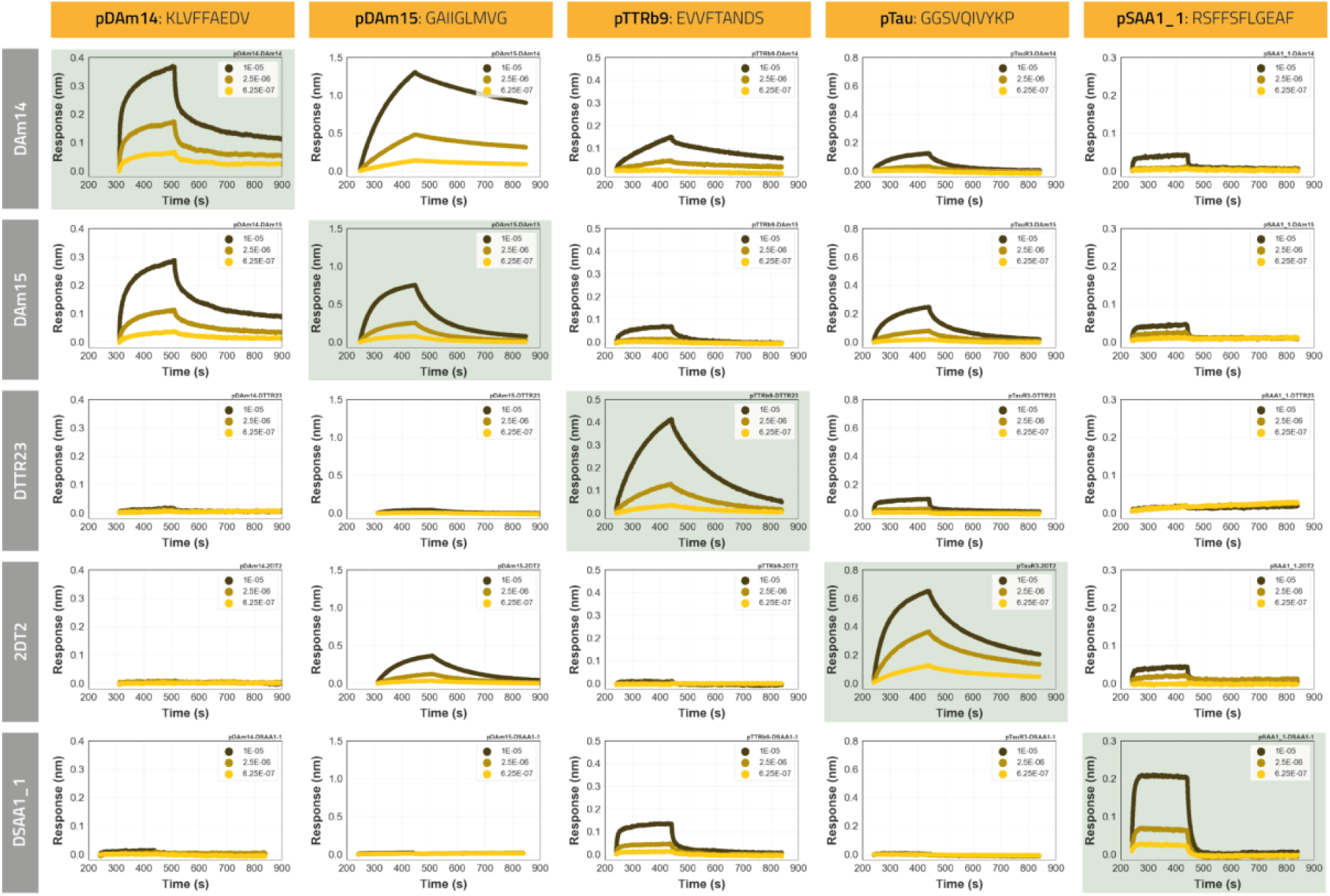
Specificity profile amyloidogenic peptide binders in BLI. Biotinylated peptides were immobilized onto octet streptavidin biosensors at equal densities and incubated with all binders in separate experiments at three concentrations (10, 2.5 and 0.625 μM). The designed on-target interactions are indicated with a light green background.

**Fig S13.**
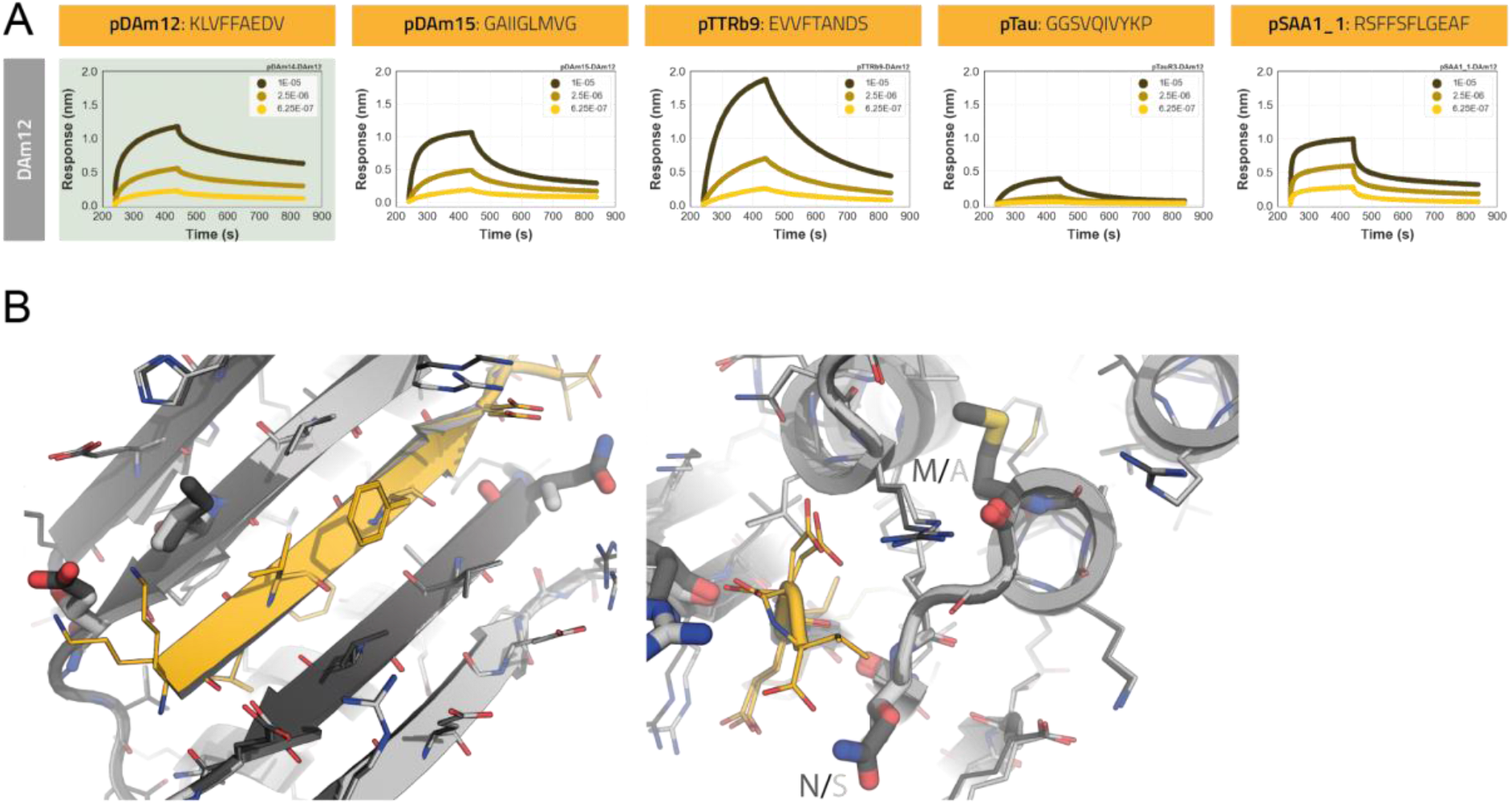
Binding characterization DAm12. **a:** DAm12 titration (10, 2.5 and 0.625 μM) against all immobilized amyloidogenic peptides in BLI. pDAm12 and pDAm14 are the same peptides. **b:** DAm12 and 14 are similar but have 4-fold difference in Aβ42 monomer binding in MDS assays. Interface close up views that highlight the interface of DAm12 (dark gray) and DAm14 (light gray). Differences in side chains are depicted in thicker sticks.

**Figure S14.**
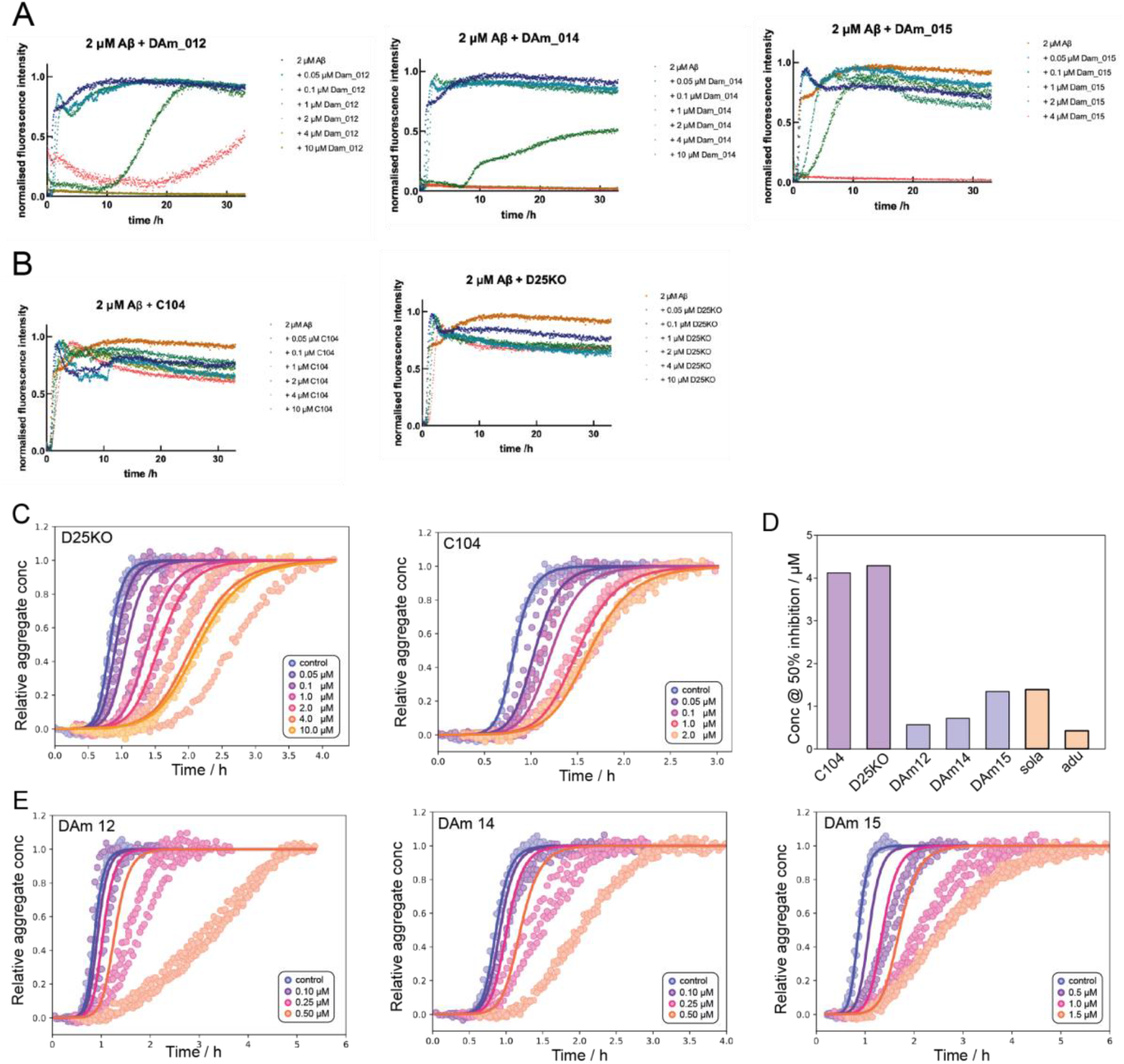
Aβ42 fibril inhibition. **a-c:** ThT Aβ42 fibril inhibition assays of the designed binders and controls that were not designed to inhibit Aβ42 aggregation. **d:** The inhibitory potential of binders, controls and clinical antibodies against Aβ42 aggregation is compared. See main fig 5d. **e:** Expected inhibitory effect due to monomer binding only. Points are ThT measurements, at a range of binder concentrations. The solid lines are produced by predicting the amount of inhibition at each binder concentration. To do so, we used the affinities of the binders to monomer to calculate the amounts of bound monomer and assumed that any monomer bound is completely removed from the aggregation reaction. Using the fits of the kinetics in the absence of binder, and the reaction orders determined previously (Cohen et al. 2013), we could then predict the expected inhibition at each binder concentration. Even for the tightest binders and assuming any bound monomer is permanently removed from the reaction, the observed inhibitory potential exceeds that expected to occur by monomer binding alone. This implies additional inhibitory mechanisms beyond interactions with monomeric Aβ42 are active.

## References

Abelein, Axel, Gefei Chen, Kristīne Kitoka, Rihards Aleksis, Filips Oleskovs, Médoune Sarr, Michael Landreh, et al. 2020. “High-Yield Production of Amyloid-β Peptide Enabled by a Customized Spider Silk Domain.” Scientific Reports 10 (1): 235.

Adams, Paul D., Pavel V. Afonine, Gábor Bunkóczi, Vincent B. Chen, Ian W. Davis, Nathaniel Echols, Jeffrey J. Headd, et al. 2010. “PHENIX: A Comprehensive Python-Based System for Macromolecular Structure Solution.” Acta Crystallographica. Section D, Biological Crystallography 66 (2): 213–21.

Alford, Rebecca F., Andrew Leaver-Fay, Jeliazko R. Jeliazkov, Matthew J. O’Meara, Frank P. DiMaio, Hahnbeom Park, Maxim V. Shapovalov, et al. 2017. “The Rosetta All-Atom Energy Function for Macromolecular Modeling and Design.” Journal of Chemical Theory and Computation 13 (6): 3031–48.

Arosio, Paolo, Thomas Müller, Luke Rajah, Emma V. Yates, Francesco A. Aprile, Yingbo Zhang, Samuel I. A. Cohen, et al. 2016. “Microfluidic Diffusion Analysis of the Sizes and Interactions of Proteins under Native Solution Conditions.” ACS Nano 10 (1): 333–41.

Bhardwaj, Gaurav, Vikram Khipple Mulligan, Christopher D. Bahl, Jason M. Gilmore, Peta J. Harvey, Olivier Cheneval, Garry W. Buchko, et al. 2016. “Accurate de Novo Design of Hyperstable Constrained Peptides.” Nature 538 (7625): 329–35.

Bloom, George S. 2014. “Amyloid-β and Tau: The Trigger and Bullet in Alzheimer Disease Pathogenesis.” JAMA Neurology 71 (4): 505–8.

Boutajangout, Allal, Hanna Lindberg, Abdulaziz Awwad, Arun Paul, Rabaa Baitalmal, Ismail Almokyad, Ingmarie Höidén-Guthenberg, et al. 2019. “Affibody-Mediated Sequestration of Amyloid β Demonstrates Preventive Efficacy in a Transgenic Alzheimer’s Disease Mouse Model.” Frontiers in Aging Neuroscience 11 (March): 64.

Boyken, Scott E., Zibo Chen, Benjamin Groves, Robert A. Langan, Gustav Oberdorfer, Alex Ford, Jason M. Gilmore, et al. 2016. “De Novo Design of Protein Homo-Oligomers with Modular Hydrogen-Bond Network-Mediated Specificity.” Science 352 (6286): 680–87.

Brunette, T. J., Fabio Parmeggiani, Po-Ssu Huang, Gira Bhabha, Damian C. Ekiert, Susan E. Tsutakawa, Greg L. Hura, John A. Tainer, and David Baker. 2015. “Exploring the Repeat Protein Universe through Computational Protein Design.” Nature 528 (7583): 580–84.

Chaudhury, Sidhartha, Sergey Lyskov, and Jeffrey J. Gray. 2010. “PyRosetta: A Script-Based Interface for Implementing Molecular Modeling Algorithms Using Rosetta.” Bioinformatics 26 (5): 689–91.

Chen, Gefei, Axel Abelein, Harriet E. Nilsson, Axel Leppert, Yuniesky Andrade-Talavera, Simone Tambaro, Lovisa Hemmingsson, et al. 2017. “Bri2 BRICHOS Client Specificity and Chaperone Activity Are Governed by Assembly State.” Nature Communications 8 (1): 2081.

Chen, Guo-Fang, Ting-Hai Xu, Yan Yan, Yu-Ren Zhou, Yi Jiang, Karsten Melcher, and H. Eric Xu. 2017. “Amyloid Beta: Structure, Biology and Structure-Based Therapeutic Development.” Acta Pharmacologica Sinica 38 (9): 1205–35.

Chiti, Fabrizio, and Christopher M. Dobson. 2006. “Protein Misfolding, Functional Amyloid, and Human Disease.” Annual Review of Biochemistry 75: 333–66.

Cohen, Samuel I. A., Sara Linse, Leila M. Luheshi, Erik Hellstrand, Duncan A. White, Luke Rajah, Daniel E. Otzen, Michele Vendruscolo, Christopher M. Dobson, and Tuomas P. J. Knowles. 2013. “Proliferation of Amyloid-β42 Aggregates Occurs through a Secondary Nucleation Mechanism.” Proceedings of the National Academy of Sciences of the United States of America 110 (24): 9758–63.

Dauparas, J., I. Anishchenko, N. Bennett, H. Bai, R. J. Ragotte, L. F. Milles, B. I. M. Wicky, et al. 2022. “Robust Deep Learning Based Protein Sequence Design Using ProteinMPNN.” bioRxiv. https://doi.org/10.1101/2022.06.03.494563.

Emsley, Paul, and Kevin Cowtan. 2004. “Coot: Model-Building Tools for Molecular Graphics.” Acta Crystallographica. Section D, Biological Crystallography 60 (Pt 12 Pt 1): 2126–32.

Fleishman, Sarel J., Andrew Leaver-Fay, Jacob E. Corn, Eva-Maria Strauch, Sagar D. Khare, Nobuyasu Koga, Justin Ashworth, et al. 2011. “RosettaScripts: A Scripting Language Interface to the Rosetta Macromolecular Modeling Suite.” PloS One 6 (6): e20161.

Gamblin, T. Chris, Feng Chen, Angara Zambrano, Aida Abraha, Sarita Lagalwar, Angela L. Guillozet, Meiling Lu, et al. 2003. “Caspase Cleavage of Tau: Linking Amyloid and Neurofibrillary Tangles in Alzheimer’s Disease.” Proceedings of the National Academy of Sciences of the United States of America 100 (17): 10032–37.

Gremer, Lothar, Daniel Schölzel, Carla Schenk, Elke Reinartz, Jörg Labahn, Raimond B. G. Ravelli, Markus Tusche, et al. 2017. “Fibril Structure of Amyloid-β(1-42) by Cryo-Electron Microscopy.” Science 358 (6359): 116–19.

Guerrero-Ferreira, Ricardo, Nicholas Mi Taylor, Daniel Mona, Philippe Ringler, Matthias E. Lauer, Roland Riek, Markus Britschgi, and Henning Stahlberg. 2018. “Cryo-EM Structure of Alpha-Synuclein Fibrils.” eLife 7 (July). https://doi.org/10.7554/eLife.36402.

Hansen, D. Flemming, Pramodh Vallurupalli, and Lewis E. Kay. 2008. “An Improved 15N Relaxation Dispersion Experiment for the Measurement of Millisecond Time-Scale Dynamics in Proteins.” The Journal of Physical Chemistry. B 112 (19): 5898–5904.

Hosseinzadeh, Parisa, Gaurav Bhardwaj, Vikram Khipple Mulligan, Matthew D. Shortridge, Timothy W. Craven, Fátima Pardo-Avila, Stephen A. Rettie, et al. 2017. “Comprehensive Computational Design of Ordered Peptide Macrocycles.” Science 358 (6369): 1461–66.

Hsia, Yang, Rubul Mout, William Sheffler, Natasha I. Edman, Ivan Vulovic, Young-Jun Park, Rachel L. Redler, et al. 2021. “Design of Multi-Scale Protein Complexes by Hierarchical Building Block Fusion.” Nature Communications 12 (1): 2294.

Huang, Po-Ssu, Yih-En Andrew Ban, Florian Richter, Ingemar Andre, Robert Vernon, William R. Schief, and David Baker. 2011. “RosettaRemodel: A Generalized Framework for Flexible Backbone Protein Design.” PloS One 6 (8): e24109.

Iakovleva, Irina, Michael Hall, Melanie Oelker, Linda Sandblad, Intissar Anan, and A. Elisabeth Sauer-Eriksson. 2021. “Structural Basis for Transthyretin Amyloid Formation in Vitreous Body of the Eye.” Nature Communications 12 (1): 7141.

Jiang, Bin, Binhan Yu, Xu Zhang, Maili Liu, and Daiwen Yang. 2015. “A (15)N CPMG Relaxation Dispersion Experiment More Resistant to Resonance Offset and Pulse Imperfection.” Journal of Magnetic Resonance 257 (August): 1–7.

Jiang, Yi Xiao, Qin Cao, Michael R. Sawaya, Romany Abskharon, Peng Ge, Michael DeTure, Dennis W. Dickson, et al. 2022. “Amyloid Fibrils in FTLD-TDP Are Composed of TMEM106B and Not TDP-43.” Nature 605 (7909): 304–9.

Joosten, Robbie P., Fei Long, Garib N. Murshudov, and Anastassis Perrakis. 2014. “The PDB_REDO Server for Macromolecular Structure Model Optimization.” IUCrJ 1 (Pt 4): 213–20.

Jumper, John, Richard Evans, Alexander Pritzel, Tim Green, Michael Figurnov, Olaf Ronneberger, Kathryn Tunyasuvunakool, et al. 2021. “Highly Accurate Protein Structure Prediction with AlphaFold.” Nature 596 (7873): 583–89.

Kabsch, Wolfgang. 2010. “XDS.” Acta Crystallographica. Section D, Biological Crystallography 66 (Pt 2): 125–32.

Knowles, Tuomas P. J., Michele Vendruscolo, and Christopher M. Dobson. 2014. “The Amyloid State and Its Association with Protein Misfolding Diseases.” Nature Reviews. Molecular Cell Biology 15 (6): 384–96.

Koepnick, Brian, Jeff Flatten, Tamir Husain, Alex Ford, Daniel-Adriano Silva, Matthew J. Bick, Aaron Bauer, et al. 2019. “De Novo Protein Design by Citizen Scientists.” Nature 570 (7761): 390–94.

Koga, Nobuyasu, Rie Tatsumi-Koga, Gaohua Liu, Rong Xiao, Thomas B. Acton, Gaetano T. Montelione, and David Baker. 2012. “Principles for Designing Ideal Protein Structures.” Nature 491 (7423): 222–27.

Kronqvist, Nina, Médoune Sarr, Anton Lindqvist, Kerstin Nordling, Martins Otikovs, Luca Venturi, Barbara Pioselli, et al. 2017. “Efficient Protein Production Inspired by How Spiders Make Silk.” Nature Communications 8 (May): 15504.

Leman, Julia Koehler, Brian D. Weitzner, Steven M. Lewis, Jared Adolf-Bryfogle, Nawsad Alam, Rebecca F. Alford, Melanie Aprahamian, et al. 2020. “Macromolecular Modeling and Design in Rosetta: Recent Methods and Frameworks.” Nature Methods 17 (7): 665–80.

Liberta, Falk, Sarah Loerch, Matthies Rennegarbe, Angelika Schierhorn, Per Westermark, Gunilla T. Westermark, Bouke P. C. Hazenberg, Nikolaus Grigorieff, Marcus Fändrich, and Matthias Schmidt. 2019. “Cryo-EM Fibril Structures from Systemic AA Amyloidosis Reveal the Species Complementarity of Pathological Amyloids.” Nature Communications 10 (1): 1104.

Linse, Sara, Tom Scheidt, Katja Bernfur, Michele Vendruscolo, Christopher M. Dobson, Samuel I. A. Cohen, Eimantas Sileikis, et al. 2020. “Kinetic Fingerprints Differentiate the Mechanisms of Action of Anti-Aβ Antibodies.” Nature Structural & Molecular Biology 27 (12): 1125–33.

Lin, Yu-Ru, Nobuyasu Koga, Rie Tatsumi-Koga, Gaohua Liu, Amanda F. Clouser, Gaetano T. Montelione, and David Baker. 2015. “Control over Overall Shape and Size in de Novo Designed Proteins.” Proceedings of the National Academy of Sciences of the United States of America 112 (40): E5478–85.

Lin, Yu-Ru, Nobuyasu Koga, Sergey M. Vorobiev, and David Baker. 2017. “Cyclic Oligomer Design with de Novo αβ-Proteins: Fixed and Flexible Backbone Cyclic Oligomer Design Using De Novo αβ Proteins.” Protein Science: A Publication of the Protein Society 26 (11): 2187–94.

Lu, Jinghua, Yadong Yu, Iowis Zhu, Yifan Cheng, and Peter D. Sun. 2014. “Structural Mechanism of Serum Amyloid A-Mediated Inflammatory Amyloidosis.” Proceedings of the National Academy of Sciences of the United States of America 111 (14): 5189–94.

Marsh, Joseph A., Vinay K. Singh, Zongchao Jia, and Julie D. Forman-Kay. 2006. “Sensitivity of Secondary Structure Propensities to Sequence Differences between Alpha- and Gamma-Synuclein: Implications for Fibrillation.” Protein Science: A Publication of the Protein Society 15 (12): 2795–2804.

McCoy, Airlie J., Ralf W. Grosse-Kunstleve, Paul D. Adams, Martyn D. Winn, Laurent C. Storoni, and Randy J. Read. 2007. “Phaser Crystallographic Software.” Journal of Applied Crystallography 40 (Pt 4): 658–74.

Meisl, Georg, Julius B. Kirkegaard, Paolo Arosio, Thomas C. T. Michaels, Michele Vendruscolo, Christopher M. Dobson, Sara Linse, and Tuomas P. J. Knowles. 2016. “Molecular Mechanisms of Protein Aggregation from Global Fitting of Kinetic Models.” Nature Protocols 11 (2): 252–72.

Muchtar, E., A. Dispenzieri, H. Magen, M. Grogan, M. Mauermann, E. D. McPhail, P. J. Kurtin, et al. 2021. “Systemic Amyloidosis from A (AA) to T (ATTR): A Review.” Journal of Internal Medicine 289 (3): 268–92.

Müller, Thomas, Paolo Arosio, Luke Rajah, Samuel I. A. Cohen, Emma V. Yates, Michele Vendruscolo, Chrisopher M. Dobson, and Tuomas P. J. Knowles. 2015. “Particle-Based Simulations of Steady-State Mass Transport at High Péclet Numbers.” arXiv [physics.flu-Dyn]. arXiv. http://arxiv.org/abs/1510.05126.

Murshudov, G. N., A. A. Vagin, and E. J. Dodson. 1997. “Refinement of Macromolecular Structures by the Maximum-Likelihood Method.” Acta Crystallographica. Section D, Biological Crystallography 53 (Pt 3): 240–55.

Otwinowski, Zbyszek, and Wladek Minor. 1997. “[20] Processing of X-Ray Diffraction Data Collected in Oscillation Mode.” In Methods in Enzymology, 276:307–26. Academic Press.

Panza, Francesco, Madia Lozupone, Giancarlo Logroscino, and Bruno P. Imbimbo. 2019. “A Critical Appraisal of Amyloid-β-Targeting Therapies for Alzheimer Disease.” Nature Reviews. Neurology 15 (2): 73–88.

Qin, Dong, Younan Xia, and George M. Whitesides. 2010. “Soft Lithography for Micro- and Nanoscale Patterning.” Nature Protocols 5 (3): 491–502.

Remaut, Han, and Gabriel Waksman. 2006. “Protein-Protein Interaction through Beta-Strand Addition.” Trends in Biochemical Sciences 31 (8): 436–44.

Sahtoe, Danny D., Adrian Coscia, Nur Mustafaoglu, Lauren M. Miller, Daniel Olal, Ivan Vulovic, Ta-Yi Yu, et al. 2021. “Transferrin Receptor Targeting by de Novo Sheet Extension.” Proceedings of the National Academy of Sciences of the United States of America 118 (17). https://doi.org/10.1073/pnas.2021569118.

Sahtoe, Danny D., Florian Praetorius, Alexis Courbet, Yang Hsia, Basile I. M. Wicky, Natasha I. Edman, Lauren M. Miller, et al. 2022. “Reconfigurable Asymmetric Protein Assemblies through Implicit Negative Design.” Science 375 (6578): eabj7662.

Sattler, Michael, Jürgen Schleucher, and Christian Griesinger. 1999. “Heteronuclear Multidimensional NMR Experiments for the Structure Determination of Proteins in Solution Employing Pulsed Field Gradients.” Progress in Nuclear Magnetic Resonance Spectroscopy 34 (2): 93–158.

Schmidt, Matthias, Sebastian Wiese, Volkan Adak, Jonas Engler, Shubhangi Agarwal, Günter Fritz, Per Westermark, Martin Zacharias, and Marcus Fändrich. 2019. “Cryo-EM Structure of a Transthyretin-Derived Amyloid Fibril from a Patient with Hereditary ATTR Amyloidosis.” Nature Communications 10 (1): 5008.

Schneider, Matthias M., Saurabh Gautam, Therese W. Herling, Ewa Andrzejewska, Georg Krainer, Alyssa M. Miller, Victoria A. Trinkaus, et al. 2021. “The Hsc70 Disaggregation Machinery Removes Monomer Units Directly from α-Synuclein Fibril Ends.” Nature Communications 12 (1): 5999.

Shammas, Sarah L., Michael D. Crabtree, Liza Dahal, Basile I. M. Wicky, and Jane Clarke. 2016. “Insights into Coupled Folding and Binding Mechanisms from Kinetic Studies.” The Journal of Biological Chemistry 291 (13): 6689–95.

Shi, Yang, Wenjuan Zhang, Yang Yang, Alexey G. Murzin, Benjamin Falcon, Abhay Kotecha, Mike van Beers, et al. 2021. “Structure-Based Classification of Tauopathies.” Nature 598 (7880): 359–63.

Stranges, P. Benjamin, Mischa Machius, Michael J. Miley, Ashutosh Tripathy, and Brian Kuhlman. 2011. “Computational Design of a Symmetric Homodimer Using β-Strand Assembly.” Proceedings of the National Academy of Sciences of the United States of America 108 (51): 20562–67.

Studier, F. William. 2005. “Protein Production by Auto-Induction in High Density Shaking Cultures.” Protein Expression and Purification 41 (1): 207–34.

Thacker, Dev, Mara Bless, Mohammad Barghouth, Enming Zhang, and Sara Linse. 2022. “A Palette of Fluorescent Aβ42 Peptides Labelled at a Range of Surface-Exposed Sites.” International Journal of Molecular Sciences 23 (3). https://doi.org/10.3390/ijms23031655.

Tsai, C. J., D. Xu, and R. Nussinov. 1998. “Protein Folding via Binding and Vice Versa.” Folding and Design 3 (4): R71–80.

Tyka, Michael D., Kenneth Jung, and David Baker. 2012. “Efficient Sampling of Protein Conformational Space Using Fast Loop Building and Batch Minimization on Highly Parallel Computers.” Journal of Computational Chemistry 33 (31): 2483–91.

Várnai, P., and T. Balla. 1998. “Visualization of Phosphoinositides That Bind Pleckstrin Homology Domains: Calcium- and Agonist-Induced Dynamic Changes and Relationship to Myo-[3H]inositol-Labeled Phosphoinositide Pools.” The Journal of Cell Biology 143 (2): 501–10.

Watkins, Andrew M., and Paramjit S. Arora. 2014. “Anatomy of β-Strands at Protein-Protein Interfaces.” ACS Chemical Biology 9 (8): 1747–54.

Wicky, B. I. M., L. F. Milles, A. Courbet, R. J. Ragotte, J. Dauparas, E. Kinfu, S. Tipps, et al. 2022. “Hallucinating Symmetric Protein Assemblies.” Science 378 (6615): 56–61.

Williams, Christopher J., Jeffrey J. Headd, Nigel W. Moriarty, Michael G. Prisant, Lizbeth L. Videau, Lindsay N. Deis, Vishal Verma, et al. 2018. “MolProbity: More and Better Reference Data for Improved All-Atom Structure Validation.” Protein Science: A Publication of the Protein Society 27 (1): 293–315.

Winn, Martyn D., Charles C. Ballard, Kevin D. Cowtan, Eleanor J. Dodson, Paul Emsley, Phil R. Evans, Ronan M. Keegan, et al. 2011. “Overview of the CCP4 Suite and Current Developments.” Acta Crystallographica. Section D, Biological Crystallography 67 (Pt 4): 235–42.

Wright, Peter E., and H. Jane Dyson. 2009. “Linking Folding and Binding.” Current Opinion in Structural Biology 19 (1): 31–38.

Zwahlen, Catherine, Pascale Legault, Sébastien J. F. Vincent, Jack Greenblatt, Robert Konrat, and Lewis E. Kay. 1997. “Methods for Measurement of Intermolecular NOEs by Multinuclear NMR Spectroscopy: Application to a Bacteriophage λ N-Peptide/boxB RNA Complex.” Journal of the American Chemical Society 119 (29): 6711–21.

